# Identification of a new pharyngeal mucosal lymphoid organ in zebrafish and other teleosts: tonsils in fish?

**DOI:** 10.1101/2023.03.13.532382

**Authors:** J Resseguier, M Nguyen-Chis, J Wohlmann, D Rigaudeau, I Salinas, SH Oehlers, GF Wiegertjes, FE Johansen, SW Qiao, EO Koppang, B Verrier, P Boudinot, G Griffiths

**Author notes:** These authors contributed equally (M Nguyen-Chis, J Wohlmann). These authors contributed equally (P Boudinot, G Griffiths).

## Abstract

The constant exposure of the fish branchial cavity to aquatic pathogens must have driven local mucosal immune responses to be extremely important for their survival. In this study, we used a universal marker for T lymphocytes/natural killer cells (ZAP70) and advanced imaging techniques to investigate the lymphoid architecture of the zebrafish branchial cavity. We identified a new lymphoid organ, which we tentatively named “Nemausean Lymphoid Organ” (NEMO), situated below the pharynx, and closely associated with gill lymphoid tissues. Besides T/NK cells, NEMO is enriched in plasma/B cells and antigen-presenting cells embedded in a network of reticulated epithelial cells. Presence of activated T cells and lymphocyte proliferation but not V(D)J recombination or hematopoiesis, suggests a function as secondary lymphoid organ. In response to infection, NEMO displays structural changes including the formation of T/NK cells clusters. NEMO and gill lymphoid aggregates form a cohesive unit within a lymphoid network that extends throughout the pharyngo-respiratory area. Collectively, our findings reveal a new mucosal lymphoid organ reminiscent of mammalian tonsils that evolved in fish. Importantly, NEMO could clearly be identified in multiple teleost fish families.

**One sentence summary:** A previously unreported lymphoid organ has been identified within the pharyngo-respiratory tract of the zebrafish, and other teleost fish, providing new insights into the immune system of teleost fish and the evolution of vertebrate immunology.

## INTRODUCTION

The survival of pluricellular organisms requires defense mechanisms against infections at barrier tissues, the interface between the environment and the host. The emergence of adaptive immunity, approximately 500 million years ago, marked a significant milestone in the defense against pathogens. Adaptive immunity is based on clonal selection of lymphocytes expressing somatically diversified genes encoding Ag receptors. The production of lymphocytes, their differentiation to naïve B or T cells, and the V(D)J recombination of genes encoding Ag receptors, occur in primary lymphoid organs. Naive lymphocytes then circulate through the bloodstream, and relocate to secondary lymphoid organs where adaptive immune responses are initiated, leading to lymphocyte activation and establishment of long-lived protective immunity. Both primary and secondary lymphoid organs are constitutive and develop at predetermined locations (*1*). During evolution, the emergence of secondary lymphoid organs has been essential in facilitating adaptive immune responses by providing an organizational framework favoring the co-localization of antigens and antigen-specific lymphocytes, which is necessary for the efficient induction of antibody-mediated responses (*2*). The evolution of lymphoid structures and adaptive immunity among Vertebrates have been studied and discussed over the last century (*3–10*). Surprisingly, despite teleost fish representing approximately half of all vertebrate species (http://www.iucnredlist.org), their immune system has received little attention which is particularly noteworthy given the importance of teleost fish as a vital food source for humans and animals (http://www.fao.org – The state of the world fisheries and aquaculture 2022).

The basic components of the immune system of teleost fish share many similarities with mammals (*11–13*). Most of the cells of the innate and adaptive immune systems characterized in mammals have also been identified in different species of fish, including granulocytes (*14*), innate lymphoid cells (*15*), T cells (*16*), B cells (*17–19*), and antigen-presenting cells such as macrophages (*20, 21*). Key molecular mechanisms involved in the detection of pathogens (*22*) and in the regulation of the immune responses (*23*) are also shared across jawed vertebrates. Teleost fish possess two known primary lymphoid organs: the thymus, which is responsible for the development and maturation of T cells. In fish, the thymus is composed of two separate lobules, one on the roof of each gill chamber. The thymus of teleost fish is separated into cortex like and medulla-like regions, which are not always well-defined (*24*). The other primary lymphoid organ is the kidney, a site where hematopoiesis occurs and where B cell precursors develop. The anterior part of the teleost fish kidney, the pronephros (also named “Head kidney”), is also a prominent site of immune activity associated with secondary lymphoid organ (*25*). However, in fish, it is the spleen that is considered as the main systemic secondary lymphoid organ (*26*). Recent studies have also suggested that adipose tissues may serve as additional secondary lymphoid structures (*27*). No lymph nodes nor tonsil equivalents have been observed in teleost fish. Although no clear counterparts of mammalian germinal centers have been identified in teleost fish, the stimulation of the fish immune system can induce the appearance of structures such as granulomas (*28*) and melano-macrophage centers (*29*).

In both fish and mammals, mucosal tissues provide an extensive surface that connects the organism with the outside world. Mucosal tissues facilitate critical functions such as nutrient absorption and gas exchange, however, such large surface area also increases exposure to pathogens. As in mammals, fish mucosae are protected by multiple “Mucosa-Associated Lymphoid tissues” (MALTs) which function in the immune surveillance of the mucosal interface (*30–32*). The main fish MALTs are located in the gut (GALT), the skin (SALT), the nostril (NALT), and the gills (GIALT) (*33–36*). In addition, recent studies have reported the existence of a mucosal associated lymphoid tissue associated with the mouth and the pharynx (*37, 38*). In mammals, the organization of MALTs is well-defined into regions where disorganized immune cells are scattered, hence forming a diffuse mucosal immune system, and into organized lymphoid aggregates such as Peyer’s patches in the gut and the Waldeyer’s ring of tonsils of the nasopharyngeal area (*39*). In contrast, the organization of the fish mucosal immune system has long been perceived as a set of scattered immune cells spread along mucosal territories (*18, 30, 40*). However, such an organization of the immune system would be difficult to reconcile with the evolutionary pressures exerted by the high concentration of microbes in aquatic environments (*41*). One might intuitively expect the organization of the fish immune system to be highly sophisticated. In fact, the absence organized lymphoid structures in teleost fish has been challenged by recent discoveries, such as the identification of the interbranchial lymphoid tissue (ILT) within the gills of Atlantic salmon (*Salmo salar*) in 2008 (*42*). This caused a severe breach within the paradigm of fish having a simple immune system, opening up the need for further investigations of the fish lymphoid organization.

In order to further characterize the mucosal lymphoid organization of teleost fish, we took advantage of the zebrafish as a research model system (*43*). It can be argued that the immune system of the zebrafish is one of the best characterized among teleost fish. It is also ideally suited for imaging, for whole-organism investigations, and benefits greatly from numerous molecular tools, including many publicly available genetically-modified strains. The zebrafish is also an established animal model to study human diseases and immune mechanisms (*11, 44–47*), as well as an excellent model system to understand mechanisms relevant for aquaculture (*48*).

In a previous study (*36*), using high-resolution 3D imaging of the zebrafish gills we characterized the organization of the gill-associated lymphoid tissue (GIALT) and identified its compartmentalization into segments where immune cells are unorganized, and two lymphoid aggregates that display features of secondary lymphoid organs: the ILT and a newly identified lymphoid tissue that we called the amphibranchial lymphoid tissue (ALT). These findings revealed a higher degree of organization of fish MALTs and support our contention that there is still much to learn about the organization of the fish immune system. This is particularly true for the branchial cavity (also named gill chambers or pharyngeal cavities), which represents one of the least understood parts of the fish anatomy despite its importance for many critical functions such as breathing and ionic homeostasis. The branchial cavity in fact represents a huge interface that is constantly accessible to water-borne microorganisms and debris. It also contains essential immune structures, including the thymus and gill lymphoid tissues. The branchial cavity consists of two chambers, one on each side of the head, that are bridged by the pharynx in the middle and are open to the outside *via* the operculum slits. The region below the pharynx that separates the gill chambers is called pharyngeal isthmus. Depending on the fish species, the gill chambers can be entirely separated from each other, connected only *via* the pharynx, as in zebrafish, or they can also merge below the pharyngeal isthmus, such as in Atlantic salmon (*49*). The whole branchial cavity is lined by a non-keratinized squamous “pharyngeal” epithelium (*8, 50*), which will be named “cavo-branchial epithelium” in the present study to distinguish it from the histologically distinct epithelium that covers the pharynx. Finally, each zebrafish gill chamber displays a set of four gill arches that connect the anterior sub-pharyngeal region to the upper area of the branchial cavity, with each gill arch bearing two ALTs and one ILT (*36*). The overall anatomy of the branchial cavity and the gills are illustrated in **Figure S1** and **Figure S2**.

In order to investigate the whole branchial cavity, we used cryosections of adult zebrafish heads in which we labeled tissue structures with fluorescent probes and then identified lymphoid structures using an antibody targeting a highly conserved epitope of the kinase ZAP70, a marker of T/NK cells. Our observations revealed a prominent lymphoid organ along the sub-pharyngeal region of the branchial cavity that to the best of our knowledge had not been previously described, and which we named the “Nemausean Lymphoid Organ” (NEMO).

## RESULTS

### Identification of the Nemausean Lymphoid organ, a new lymphoid structure inside the branchial cavity

As the zebrafish branchial cavity constitutes a complex anatomical territory, in order to investigate this region in its entirety we conducted high-resolution 3D multi-fields of view imaging of whole branchial cavity cryosections (30 µm) from adults. These sections were stained with fluorescent phalloidin to label F-actin and DAPI to stain DNA, facilitating the identification of tissue structures. We focused on the immunolabeling of “Zeta-chain-associated protein kinase 70 (ZAP70), a T cell / Natural Killer (NK) cell marker (*51*), to reveal the organization of lymphoid tissues. The anti-ZAP70 antibody was used extensively in our earlier studies, and has a good affinity across many species (*36, 52*). Additional evidences of its specificity are presented **Fig.S3**.

During our initial exploration of the lower region of the branchial cavity using cryosections of healthy adult wild-type (wt) zebrafish at various orientations (Fish N=10) (**Fig.1 A**), we found a previously undescribed mucosal lymphoid organ below the pharynx, at the convergence of the gill arches with the sub-pharyngeal isthmus. We tentatively named it “Nemausean Lymphoid Organ” (NEMO) inspired by the Gallic-Roman “Nemausus - Nemausicae” mythology associated with protection, water, and healing. NEMO was present in all analyzed fish.

**Figure 1.**
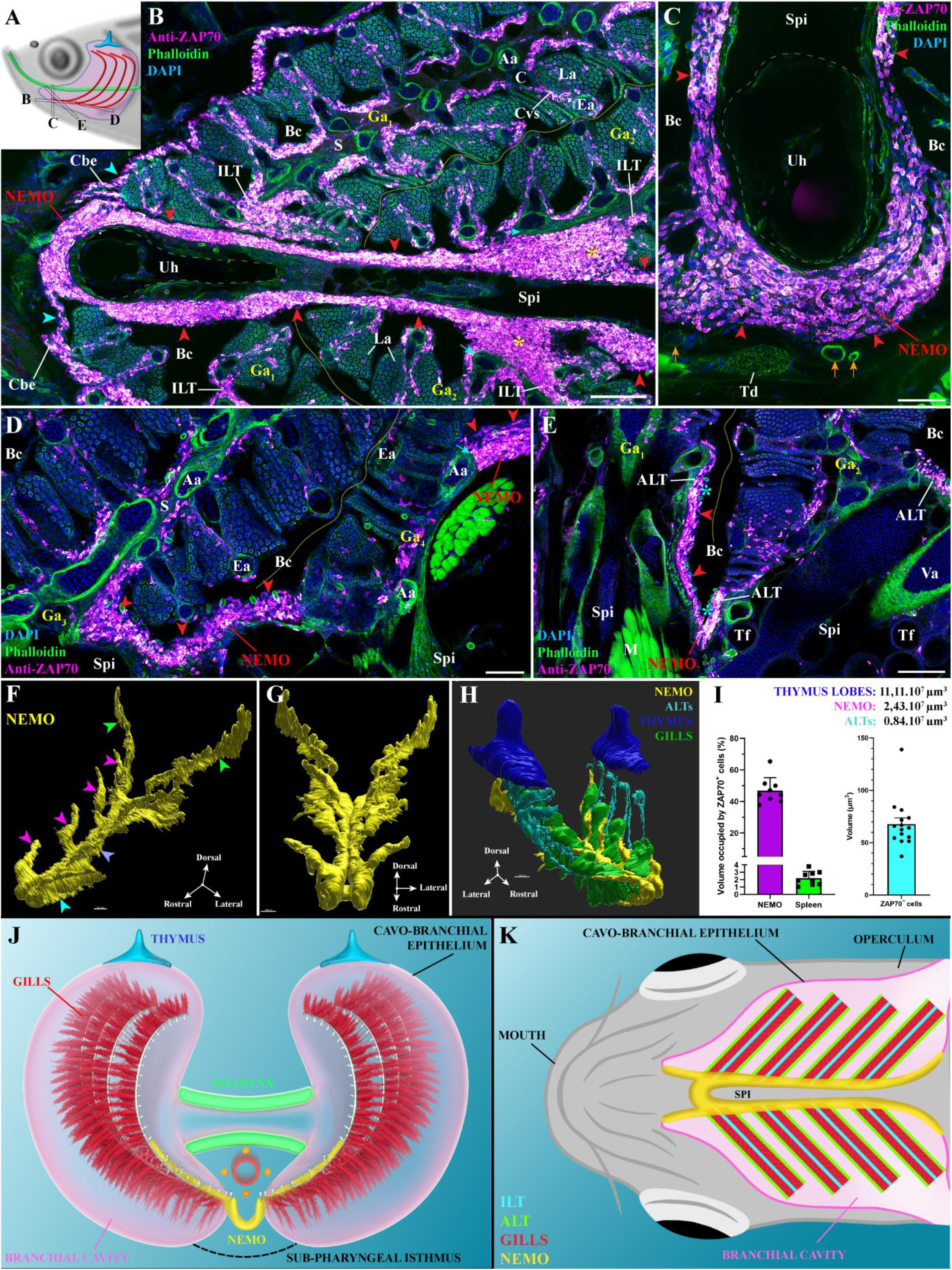
General organization and localization of the adult zebrafish NEMO. (A) Scheme illustrating the different orientations of the adult zebrafish NEMO images acquired from 30 μm whole-head cryosections, and which highlight the position of the thymus (blue), pharynx (green), and gills (red). (B-E). NEMO (red arrowheads) wraps around the urohyal bone (B,C), and extends along the sub-pharyngeal isthmus toward the posterior end of the gill chambers (D). NEMO is connected to the interbranchial lymphoid tissues (B - yellow stars), the amphibranchial lymphoid tissues (E – cyan stars), and the cavobranchial epithelium (A - cyan arrowheads). NEMO is in close proximity to gills afferent arteries (B – cyan arrows) and other endothelial vessels (C - orange arrows). (F-G) NEMO 3D reconstruction obtained with serial confocal tomography of a ZAP70-labeled wholemount head of zebrafish (15 wpf), revealing a segmentation into 4 anatomic regions: the front end wrapped around the urohyal bone (cyan arrowhead), antler-like protrusions (magenta arrowheads), the core (blue arrow), and the posterior end (green arrowheads). (H) 3D reconstruction of NEMO (yellow), ALTs (cyan), thymus lobes (blue) and the ventral end of gill arches (green). (I) Volumes of different lymphoid structures from the 3D reconstruction, average volume occupied by T/NK cells in NEMO and the spleen, and average volume of a single T/NK cell. (J,K) Simplified illustrations of NEMO’s localization within the branchial cavity, as observed from the front (J) or from below (K). Illustrations made by Ella maru studio and K.Zulkefli. Annotations: Aa, Afferent artery; ALT, Amphibranchial lymphoid tissue; Bc, Branchial cavity; C, Cartilage; Cbe, Cavo-branchial epithelium; Cvs, Central venous sinus; Ea, Efferent artery; Ga, Gill arch; ILT, Interbranchial lymphoid tissue; La, Lamellae; M, Muscles; NEMO, Nemausean lymphoid organ; S, Septum; Spi, Sub-pharyngeal isthmus; Td, Tendon; Tf, Thyroid follicle; Uh, Urohyal bone and Va, Ventral aorta. Scale bars: 150 μm (H), 100 μm (B, E-G), 50 μm (D), and 40 μm (C).

Analysis of the cryosections revealed that NEMO constitutes a large structure enriched in ZAP70-positive cells located within the squamous mucosal epithelium lining the sub pharyngeal isthmus, a region under the pharynx that separates the two gill chambers (**Fig.1 B-E** – red arrowheads). NEMO wraps around the urohyal bone at the anterior end of the branchial cavity (**Fig.1 B,C**), joining the two gill chambers, before extending along each sides of the sub pharyngeal isthmus until it reaches the posterior end of the branchial cavity (**Fig.1 D**). Along its length, NEMO is connected to all the twenty-four gill lymphoid aggregates: the eight ILTs (**Fig.1 B** – yellow stars) and sixteen ALTs (**Fig.1 E** – cyan stars). Intriguingly, we could not define any clear separation between ILT/ALT and NEMO at these connection sites, suggesting that lymphoid structures of the branchial cavity may function as a single integrated unit. This could have important implications regarding the role of the branchial cavity for the fish immunity and its development. Other images displaying NEMO and its direct connection with gill lymphoid aggregates are shown in **Fig.S4 A-D**. In addition, the expression of the kinase Lck gene, another T cell marker, was investigated using *Tg(lck:EGFP)*(*16*) transgenic adult zebrafish line and confirmed the presence of a large reservoir of T/NK cells along the sub pharyngeal region of the branchial cavity (**Fig.S4 E,F)**.

### 3D structure of NEMO

These data indicated that NEMO exhibits a sophisticated architecture that is difficult to fully capture from cryosections. We therefore built a more accurate reconstruction of NEMO’s structure in three-dimensions by adapting our imaging approach from cryosections to whole young adult wild-type zebrafish heads (15 weeks post-fertilization (wpf)) to reveal the distribution of T/NK cells within the whole branchial cavity area using serial confocal tomography. This automated technique combines sectioning with a vibratome and imaging with a confocal microscope using a robot. This approach enabled the assembly of a NEMO 3D structure from over 700 imaged layers, thereby defining NEMO boundaries using the distribution of the ZAP70 signal (**Fig.1 F,G**). This accurate representation revealed the segmentation of NEMO into four distinct anatomical sub-regions: 1. The anterior-most region that wraps around the urohyal bone (**Fig.1 F** – cyan arrowhead) 2. The four “antler-like” protrusions that each connect with two ALTs (**Fig.1 F** – magenta arrowheads) 3. The core that extends along the sub-pharyngeal isthmus (**Fig.1 F** – blue arrowhead), and 4. The posterior end that starts after the 4^th^ set of gill arches and extends toward the operculum opening (**Fig.1 F** – green arrowheads). Further research will be required to determine the degree of tissue homogeneity between these four segments. The 3D reconstruction of the thymus lobes (blue), the sixteen ALTs (cyan), and the ventral extremity of the eight gill arches (green) allowed us to interpret NEMO (yellow/magenta) in the spatial context of the fish, and in particular of the branchial cavity (**Fig.1 H, Video.S1-S4**). This approach illustrated clearly the localization of NEMO along the ventral axis of the fish head. Furthermore, this global reconstruction further highlighted the continuity between NEMO antler-like protrusions with all ALTs. The localization of NEMO within the branchial cavity is further illustrated **Fig.1 J,K**.

We then estimated the volume of NEMO based on the 3D reconstruction shown in **Fig.1**. In this 15 wpf zebrafish NEMO had a volume of 2.4×10^7^ µm^3^, which is smaller than the thymus (11.1×10^7^ µm^3^; 5.5×10^7^ µm^3^ and 5.6×10^7^ µm^3^ for each lobe); at this stage, the thymus has just started to involute. Using a stereology approach and 3D reconstruction, we estimated that T/NK cells occupy 46,8% of NEMO’s volume (Sections: N=9 obtained from 3 fishes) for an average volume of 67,7 µm^3^ per ZAP70-positive cells (3D reconstructed cells: N=15 obtained from 3 fishes) (**Fig.1 I**). In this 15 wpf zebrafish, NEMO would then contain around 165 000 T/NK cells. In comparison, the number of spleen T/NK cells, assessed using a similar approach (Sections: N=9 obtained from 3 fishes) and a rough estimation of the spleen volume, was around 120 000 T/NK cells (2,2% of the spleen’s volume). Noteworthy, the overall volume of the sixteen ALTs was estimated to be 0.84×10^7^ µm^3^, therefore indicating that the ALTs represent much smaller structures compared to NEMO. Collectively, these data indicate that NEMO constitutes a prominent structure of the branchial cavity, and provide a first line of evidence that it constitutes a separate organ. Furthermore, the central position of NEMO along the two branchial cavity and at their junction suggest an ideal localization for NEMO to be an immune site centralizing immune functions protecting the branchial cavity.

### NEMO is a mucosal lymphoid organ

The next objective was to determine whether the structural organization of NEMO at the cellular level is consistent with the known organization of lymphoid organs and tissues. A hallmark of structured lymphoid aggregates, such as the thymus, lymph nodes, Peyer’s patches or tonsils in mammals is the characteristic arrangement of the immune cells within a meshwork of reticulated epithelial cells that acts as an immuno-platform (*53, 54*). This feature was a key element in classifying teleost fish ILT and ALT as lymphoid tissues (*36, 55*). We therefore labeled cryosections of NEMO with a commonly used cocktail of antibodies to reveal cytokeratins, which are essential constituents of the reticulated epithelial cell cytoskeleton. A complex network of reticulated epithelial cells was found at the boundaries (red arrowheads) and within (yellow arrowheads) the anterior segment of NEMO (**Fig.2 A**). Further analysis confirmed that this network of reticulated epithelial cells extended throughout NEMO in its entirety (**Fig.S5 A,B**). 3D reconstruction of the cytokeratin signal revealed that the arrangement of reticulated epithelial constitutes organized pockets of cells that are typical of lymphoid aggregates (**Fig.2 B** – **Video.S5**). Noteworthy, NEMO reticulated epithelial cells display a low expression of the MHC-class II gene mhc2dab (**Fig.S5 C-F**) and are connected with each other’s by hemi-desmosome (**Fig.S5 G** – cyan arrow). Both these features are shared by the reticulated epithelial cells of mammalian lymphoid aggregates (*53, 56*).

**Figure 2.**
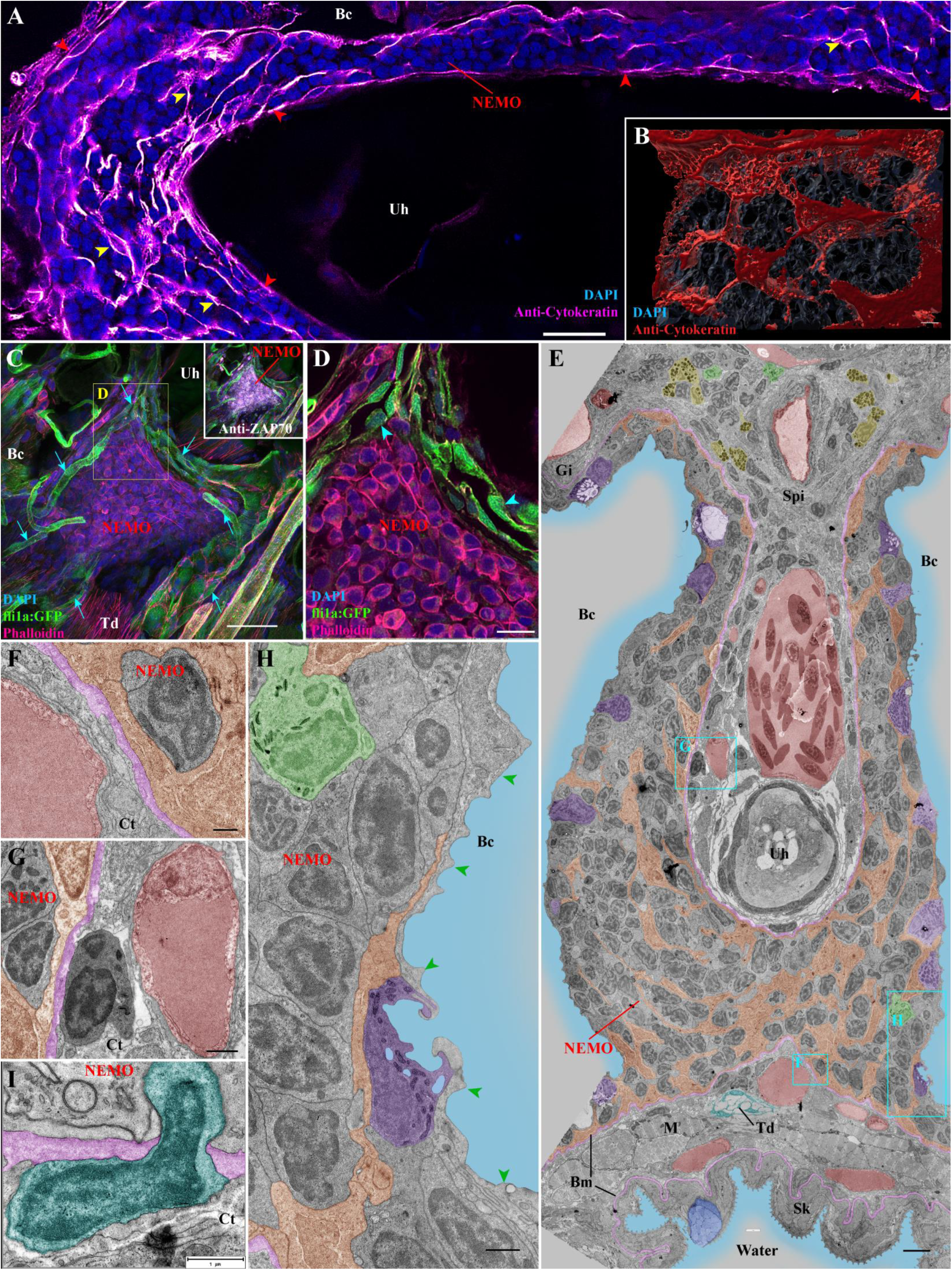
Detailed structural organization of the adult zebrafish NEMO. (A) NEMO cryosections labeled with anti-cytokeratin antibodies (magenta hot) revealing a network of reticulated epithelial cells within (yellow arrowheads) and bordering NEMO (red arrowheads). (B) 3D reconstruction illustrating the network of reticulated epithelial cells in red. (C) 3D imaging of NEMO cryosections from fli:GFP zebrafish, in which endothelial vessels are fluorescent (green). Numerous vessels are wrapped around NEMO (cyan arrows). (D) Optical section from (C) highlighting cuboidal-shaped endothelial cells (cyan arrowheads). (E-H) Ultrastructure map of a 9 wpf zebrafish NEMO transversally sectioned at the urohyal bone acquired by transmission electron microscopy. Several structures have been highlighted: Reticulated epithelial cells (orange), mucous cells (dark blue), water (light blue), ionocytes (purple), endothelial vessels (burgundy red), basement membrane (pink), neutrophils (green), basophils/mast cells (yellow), tenocytes (dark blue-green), and pavement cells (green arrowheads). (F-H) represent zoomed area from (E). (I) Cell (dark blue-green) observed across the basement membrane (pink) separating NEMO from the surrounding connective tissue. Annotations: Bc, Branchial cavity; Bm, Basement membrane; Gi, Gills; M, Muscles; Sk, Skin; Spi, Sub-pharyngeal isthmus; Td, Tendon and Uh, Urohyal bone. Scale bars: 30 μm (C), 20 μm (A), 10 μm (D), 4 μm (E), 3 μm (B), 1 μm (G,H,I), and 500 nm (F).

In order to sustain their functions, lymphoid organs require access to oxygen and nutrient supply, as well as mechanisms to facilitate immune cell trafficking. The localization of NEMO in close proximity to inhaled water ensures a continuous oxygen supply. The circulatory system functions as a conduit for nutrient delivery and immune cell trafficking throughout the body. The next question was to determine if NEMO is vascularized using 3D imaging of cryosections from an adult zebrafish line in which both vascular and lymphatic endothelial cells express a fluorescent protein (*Tg(fli1a:EGFP)* (*57*)). Although no endothelial structures were found within NEMO, numerous endothelial vessels were observed surrounding it (**Fig.2 C** – cyan arrows, **Video.S6**). This finding is consistent with the phalloidin staining, which strongly labels the smooth muscles surrounding blood vessels, observed in **Fig.1 C** (orange arrows). Opening up the 3D stacks to look at the optical sections, we found that some of the narrow vessels surrounding NEMO were lined by cuboidal endothelial cells (**Fig.2 D** – cyan arrowheads), which sometimes line fish arteries and heart endocardium (*58*). However, we could not detect around these particular vessels the characteristic layer of smooth muscles that usually surrounds fish arteries. In humans, cuboidal endothelial cells are a hallmark of the high-endothelial venules that are characteristic of mammalian lymph nodes and tonsils (*59*). Further studies will have to determine if the vessels wrapping around NEMO constitute blood vessels, conventional lymphatic vessels or the non-conventional blood/lymphatic “fine” vessels that were first reported in cod almost a century ago by Burne (*60*). In addition to the previously described vessels, NEMO also benefit from a close proximity with the prominent gill vasculature at its convergence with the gill lymphoid aggregates (**Fig.1 B,D** – cyan arrows).

### Ultrastructure of NEMO

In order to continue our investigation of NEMO at a higher resolution we used transmission electron microscopy (TEM) of ultrathin sections and a new data browsing method developed by Jens Wohlmann (manuscript in preparation) to easily access ultrastructural images at different scales over a large area. This method overcomes the challenge of analyzing complex ultrastructure of tissues at different magnifications, especially when one needs to switch from low to high-magnification in a smooth manner. Such a dynamic ultrastructure map allows the user to efficiently navigate within the biological sample. Using this approach, we assembled a detailed map (>1400 micrographs) covering a significant portion of NEMO’s anterior segment in a 9 weeks post-fertilization juvenile zebrafish (**Fig.2 E**).

The EM data highlighted a number of striking features and provided additional insights into NEMO. The network of reticulated epithelial cells was prominent (orange) and the nuclei of these cells were much less electron-dense and more elongated than the nuclei of the neighboring cells (**Fig.2 E, Fig.S5 G**). EM analysis confirmed the close proximity of NEMO to neighboring endothelial vessels (burgundy red), which were mostly separated by a thin basement membrane (pink) that forms a boundary between NEMO and the surrounding connective tissue (**Fig.2 F-G**). These observations support our view that NEMO constitutes a distinguishable entity. The ultrastructure map shows unequivocally that NEMO is only separated from the outside environment by a single layer of epithelial cells (**Fig.2 H**). These cells were predominantly pavement cells, which can be identified by their elongated shape and characteristic actin microridges (green arrowheads). Interspersed between pavement cells were mitochondria-rich cells (ionocytes) (**Fig.2 E,H** - purple). This squamous mucosal epithelium is reminiscent of the epithelium that lines the gills (*61*), which suggests they may share a similar developmental origin. The EM analysis also revealed the presence of cells that have penetrated the basement membrane bordering NEMO (**Fig.2 I** – cyan), indicating the existence of a cell traffic in or out of NEMO.

Intriguingly, whereas the thymus lobes are already present in 3 days post-fertilization zebrafish (*62*), we could not detect NEMO in 3 wpf zebrafish (not shown). Moreover in the 9 wpf juveniles we used in our ultrastructure investigation, NEMO was present but we failed to detect ILT or ALT. Looking at a publicly available atlas of zebrafish paraffin section stained with H&E (https://bio-atlas.psu.edu/zf/progress.php), we could identify structures reminiscent of NEMO in 6-7 wpf zebrafish, suggesting NEMO ontogeny would start between the 4^th^ and the 6^th^ week of development.

Collectively, our findings from both light and electron microscopy lead us to propose that NEMO is a new constitutive mucosal lymphoid organ in fish that is associated to the branchial cavity.

### NEMO is a lymphoid organ highly enriched with both B and T cells

Based on a T/NK cells marker (ZAP70), we defined NEMO as a mucosal lymphoid organ and described its location in the gill chamber area, the obvious next question was: What role does it play in the fish immune system? In order to address this question we first investigated the diversity of its immune cell populations in adult zebrafish. Our electron microscopy analysis revealed a small number of neutrophils, identified by their typical elongated granules (**Fig.2 E,H** - green). Their presence was confirmed by confocal microscopy using zebrafish in which neutrophils express fluorescent proteins (*Tg(mpx:GFP)* (*63*)) (**Fig.3 A**). Also by TEM, fish basophils/mast cells, which displayed characteristic large spherical electron-dense granules, were evident within the connective tissues adjacent to NEMO; however we did not observe any of these cells or eosinophils within NEMO itself (**Fig.2 E** –yellow).

**Figure 3.**
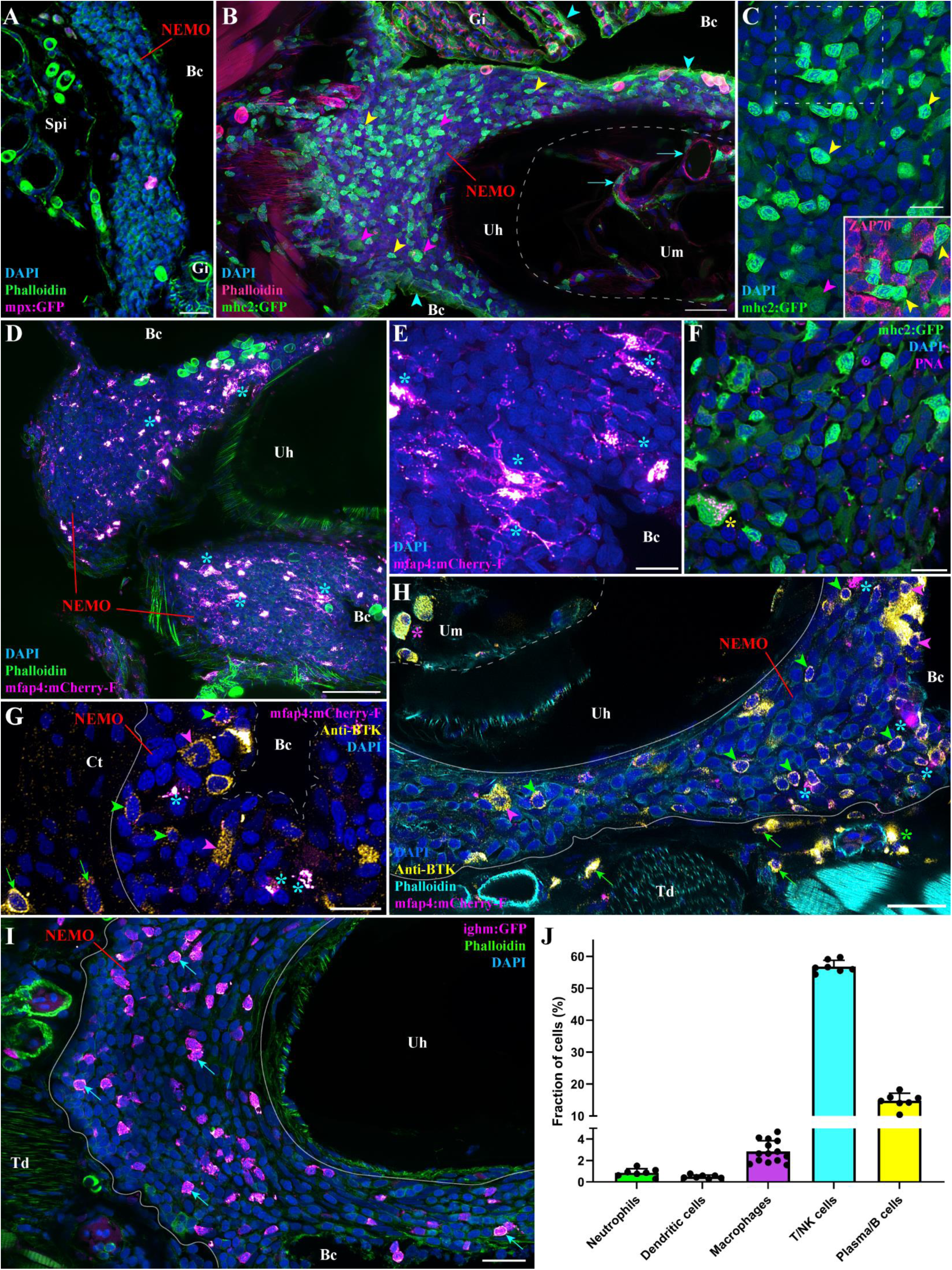
NEMO immune cell population. (A) Adult mpx:GFP zebrafish NEMO cryosection, in which neutrophils are fluorescent (Magenta hot). (B) Adult mhc2:GFP zebrafish cryosection, in which mhc2-expressing cells are fluorescent (green). Expression of MHC2 is observed in the epithelial cells (cyan arrowheads), large cells inside NEMO (magenta arrowheads), and within the marrow of the urohyal bone (cyan arrows). (C) Zoom inside NEMO of a mhc2:GFP fish highlighting large (magenta arrowheads) and small (yellow arrowheads) positive cells. The latter being negative for anti-ZAP70 labeling (cherry). (D,E) Adult mfap4:mCherry-F zebrafish NEMO cryosections, in which macrophages are fluorescent (magenta hot – cyan stars). (F) Cryosections of adult mhc2:GFP zebrafish NEMO stained with peanut agglutinin lectin (magenta hot) to reveal dendritic cell (yellow star). (G-H) Anti-BTK labeling (yellow hot) of mfap4:mCherry-F adult zebrafish cryosections revealed macrophages (magenta hot – cyan stars) as well as both BTK-positive B cells (green arrowheads) and plasma cells (magenta arrowheads) in NEMO. BTK-positive cells were also found in the connective tissue surrounding NEMO (green arrows), around endothelial structures (green stars) and within the marrow of the urohyal bone (magenta star). (I) The presence of B cells in NEMO was confirmed using ighm:GFP zebrafish, in which a subtype of B cells that express IgM is fluorescent (magenta hot – cyan arrows). (J) Quantification of the different immune cell population found in NEMO counted from at least 7 single-cell layers images originating from at least 3 fish. Annotations: Bc, Branchial cavity; Gi, Gills; Spi, Sub-pharyngeal isthmus; Td, Tendon; Uh, Urohyal bone and Um, Urohyal marrow. Scale bars: 50 μm (D), 30 μm (B), 20 μm (A, H-I), and 10 μm (C, E-G).

We next investigated the presence of antigen-presenting cells in NEMO using *Tg(mhc2dab:GFP)* (*64*) adult zebrafish (**Fig.3 B,C**). Consistent with previous studies (*36, 64*), we observed expression of the transgene in epithelial cells (cyan arrowheads), including the mhc2^low^ reticulated epithelial cells. Noteworthy, mhc2-expressing cells associated to endothelial structures were observed within the marrow of the urohyal bone (cyan arrows). Within NEMO, the presence of large mhc2^+^ cells (magenta arrowheads) suggested the presence of macrophages and/or dendritic cells, which are so-called professional antigen-presenting cells. Imaging of *Tg(mfap4:mCherry-F)* zebrafish (*65*), a zebrafish line in which macrophages express a farnesylated membrane-associated fluorescent protein, revealed many large fluorescent macrophages (**Fig.3 D,E** – cyan stars). We then labeled dendritic cells using a fluorescent peanut-agglutinin lectin, as described in (*36, 66*) (**Fig.3 E** – yellow star). This marker reveals a subset of large mhc2+ cells with a striking labeling of cytoplasmic vesicles (*66*). In addition to the large mhc2 cells, we also observed numerous small lymphocyte-like cells in NEMO that strongly expressed mhc2 but were negative for ZAP70 (**Fig.3 B,C** – yellow arrowheads).

Since B cells are usually known to lack ZAP70 and to express high amount of mhc2 proteins (*51, 67*), we hypothesized that these cells could belong to the B cell lineage. Similar to our approach using a conserved epitope of ZAP70 to label T/NK cells, we then used a well characterized antibody against human Bruton Tyrosine Kinase (BTK), an essential constituent of the B cell lineage (**Fig.3 G,H**) (*68, 69*). Since in humans BTK has also been shown to localize in subsets of macrophages (*70*), we used this antibody on mfap4:mCherry-F zebrafish in order to distinguish between macrophages and the B-cell lineage. The anti-BTK labeling revealed the presence of many B-cells (green arrowheads) and plasma cells (magenta arrowheads) within NEMO parenchyma, the adjacent connective tissue (green arrows), as well as in the marrow of the urohyal bone (magenta star). Within the surrounding connective tissue, B cells and plasma cells were often associated to endothelial structures (green star). Additional data on the anti BTK labeling are available **Fig.S6**, including the expression of mhc2 by small BTK-positive cells. The identification of B-cells in NEMO was confirmed by using the *Tg(Cau.Ighv-ighm:EGFP)* transgenic line (**Fig.3 I** – cyan arrows) (*71*), in which a subset of B-cells expressing IgM was selectively marked. Quantification of the different immune cell types in NEMO is shown in **Fig.3 J** and confirmed the predominance of lymphoid cells in NEMO: T/NK cells (56,8% of total cells), B cells (14,7%), whereas neutrophils, macrophages and dendritic cells accounted for less than 5% of the total. The remaining 24% includes the reticulated epithelial cells and the cells forming the squamous epithelial layer. Collectively, these data support our contention that NEMO has characteristics predicted for a mucosal secondary lymphoid organ. We emphasize that our analysis does not address all the cell types and cell subsets present in NEMO. For this, additional molecular analyses, such as transcriptomics, are needed.

### Functional characterization of NEMO

In order to further investigate the role of NEMO, we carried out an initial set of experiments to address whether or not NEMO could be a primary lymphoid organ. We first determined cell proliferation within NEMO using a labeling targeting the “proliferating cell nuclear antigen” (PCNA), a protein that is selectively expressed by cells engaged in cell division. As evident in **Fig.4 A** (yellow arrows), cell proliferation was prominent within NEMO. Many of the PCNA-labeled cells were also positive for ZAP70 (**Fig.4 B,C** – cyan arrows). Consistent with this result, we observed T/NK cells that displayed mitotic figures **(Fig.4 D,E – magenta arrows)**. By itself, the presence of T/NK cells proliferation is insufficient to distinguish between secondary and primary lymphoid organs. Therefore, we investigated the presence of mechanisms involved in the differentiation of lymphoid cells, which are hallmarks of primary lymphoid organs such as the thymus and kidney in zebrafish.

**Figure 4.**
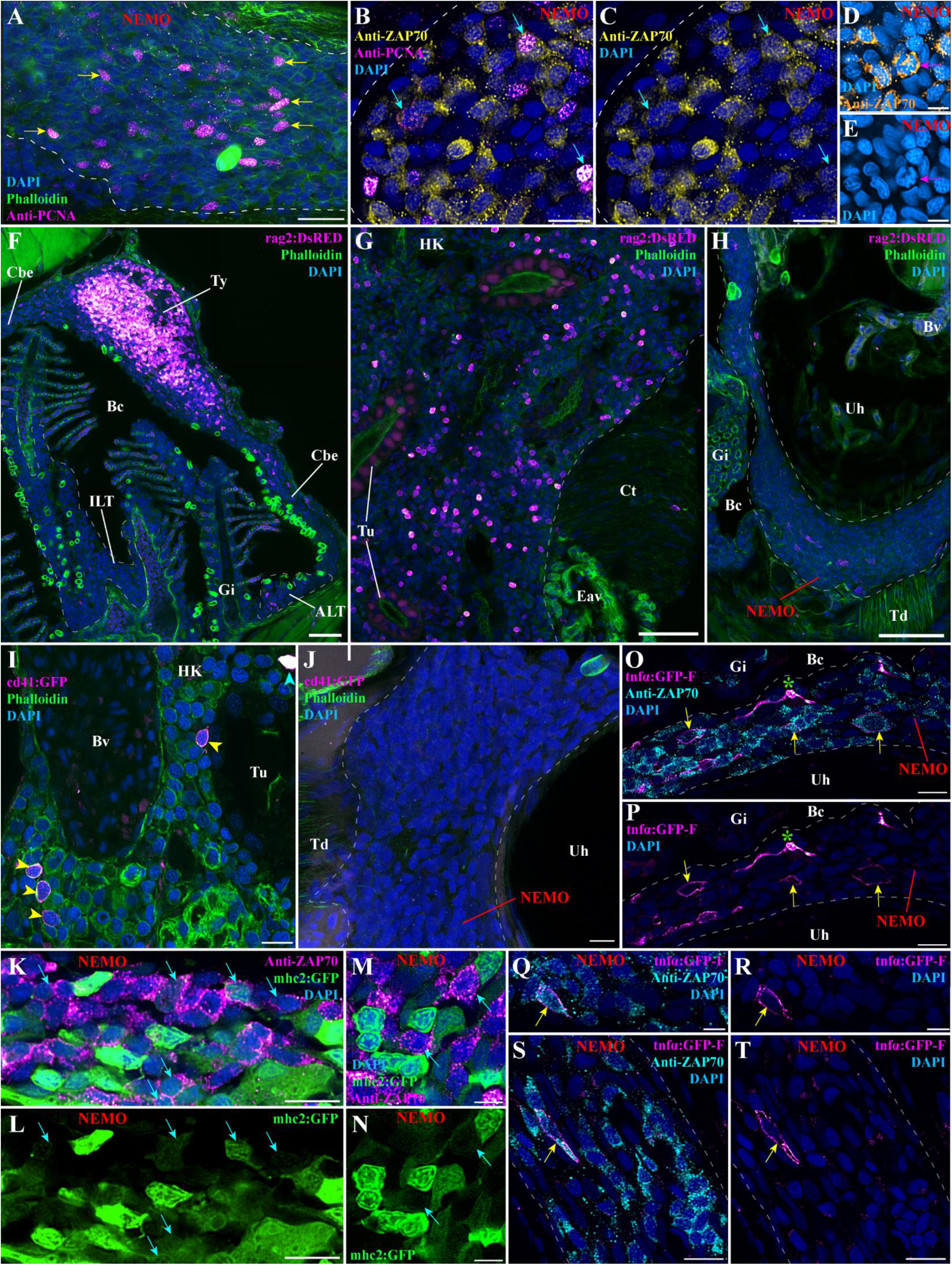
Investigation of immune function molecular markers in NEMO. (A) NEMO cryosections labeled with anti-PCNA antibody (magenta hot) to reveal cell proliferation (yellow arrows). (B-C) Cryosection co-labeled with anti-PCNA (magenta hot) and anti-ZAP70 (yellow hot) to reveal the presence of proliferative T/NK cells in NEMO (cyan arrows). (D-E) The presence of proliferative T/NK cells in NEMO was confirmed by the presence of ZAP70-positive cells (orange hot) displaying mitotic figures (magenta arrow). (F-H) Cryosections from rag2:DsRED zebrafish in which cells undergoing V(D)J recombination are fluorescent (magenta hot). Whereas numerous positive cells are found in the thymus (F) and the head-kidney (G), which are known sites of V(D)J recombination for T and B cells, almost none were observed in NEMO (H), ILTs and ALTs (F). (I,J) Cryosections from cd41:GFP zebrafish, in which thrombocyte (cyan arrowhead) are brightly fluorescent and hematopoietic stem cells are faintly fluorescent (yellow arrowhead) (magenta hot). In contrast to the expected localization of hematopoietic stem cells in the kidney (I), none were observed in NEMO (J). (K-N) Cryosections from mhc2:GFP zebrafish (green) labeled with anti-ZAP70 (magenta hot) revealed the presence of mhc2-expressing T/NK cells (cyan arrows), a feature of activated T/NK cells. (O-T) Cryosections from tnfα:GFP zebrafish NEMO, in which cells expressing the immune effector molecule TNF-α are fluorescent (magenta hot), labeled with anti-ZAP70 (cyan). Annotations: ALT, Amphibranchial lymphoid tissue; Bc, Branchial cavity; Bv, Blood vessel; Cbe, Cavobranchial epithelium; Ct, Connective tissue; Eav, Endothelium anastomotic vessels; Gi, Gills; HK, Head kidney; ILT, Interbranchial lymphoid tissue; Td, Tendon; Tu, Tubule; Ty, Thymus and Uh, Urohyal bone. Scale bars: 50 μm (F-H), 20 μm (A), 10 μm (B-C, I-L, O-P, S-T), and 5 μm (D-E, M-N, Q-R).

The protein RAG2 is an enzyme required for V(D)J recombination that is expressed by developing T cells in the thymus, and by developing B cells in the fish kidney. In order to determine if NEMO is involved in lymphocyte development we used a zebrafish line with a fluorescent reporter for rag2 expression (*Tg(rag2:DsRED*) (*72*)). In contrast to the typical high expression of rag2 found in the thymus and the head-kidney, NEMO, like the ALT and ILT (*36, 73*),did not show any significant expression of *rag2* (**Fig.4 F-H**). This result argues that NEMO does not have the primary lymphoid functions involved in B and T cell development, nor is it an additional thymus.

In adult fish the production of immune cells by hematopoiesis occurs in the kidney. This process involves hematopoietic stem cells that reside in the immune compartment of the kidney located in-between the nephrons. In zebrafish these cells are identifiable by their low expression of the protein CD41, while it is highly expressed by thrombocytes (fish analogue of platelets) (*74*). In order to determine if NEMO represents an additional site of hematopoiesis, we used the transgenic zebrafish line *Tg(cd41:GFP)*. Whereas the hematopoietic stem cells are evident within the kidney (**Fig.4 I** – yellow arrowheads), we could not observe them in NEMO (**Fig.4 J**). Collectively, our data shows that NEMO is neither involved in lymphocyte V(D)J recombination, nor in hematopoiesis, which constitutes a strong evidence that it is not a primary lymphoid organ.

We next checked whether NEMO displayed features that are characteristic of lymphoid organs involved in immune responses. During our investigation using *Tg(mhc2dab:GFP)* zebrafish we found ZAP70-positive cells that were also MHC2-positive (**Fig.4 K-N** – cyan arrows); this likely indicates the presence of activated T/NK cells in NEMO (*75, 76*). In addition, we also observed ZAP70-positive cells expressing the effector molecule TNFα (*77*), using the zebrafish line *Tg(tnfα:eGFP-F)* (*78*) (**Fig.4 O-T** – yellow arrows). Altogether, these results support the concept of NEMO being a mucosal secondary lymphoid organ.

### Structural changes in NEMO in response to viral and parasitic infection

If NEMO is indeed a mucosal secondary lymphoid organ it would be expected to be involved in immune responses to infections. Toward this goal, we first investigated zebrafish (N=3) that were naturally co-infected in a zebrafish facility with three different parasites (*Pseudoloma neurophilia*, *Pseudocapillaria tomentosa*, and *Myxidium streisingeri*) that respectively infect the nervous system, the intestines, and the kidneys (*79*). In contrast to uninfected fish (**Fig.1, Fig.S4**), the distribution of ZAP-positive cells appeared more heterogeneous and some of the labelled cells formed small local clusters (cyan stars) (**Fig.5 A,B**). Noteworthy, we could also observe an abundance of BTK-positive cells corresponding to B cells and plasma cells within the connective tissue and associated to narrow endothelial vessels of the sub-pharyngeal isthmus (**Fig.S7**). These changes gave the first hint of a structural rearrangement of NEMO in response to long-term parasitic infection. Furthermore, as none of these parasites directly infects the branchial cavity, it also reveals that NEMO’s involvement in immune response is not restricted to the branchial cavity but likely play a broader function in the overall defense of the organism.

**Figure 5.**
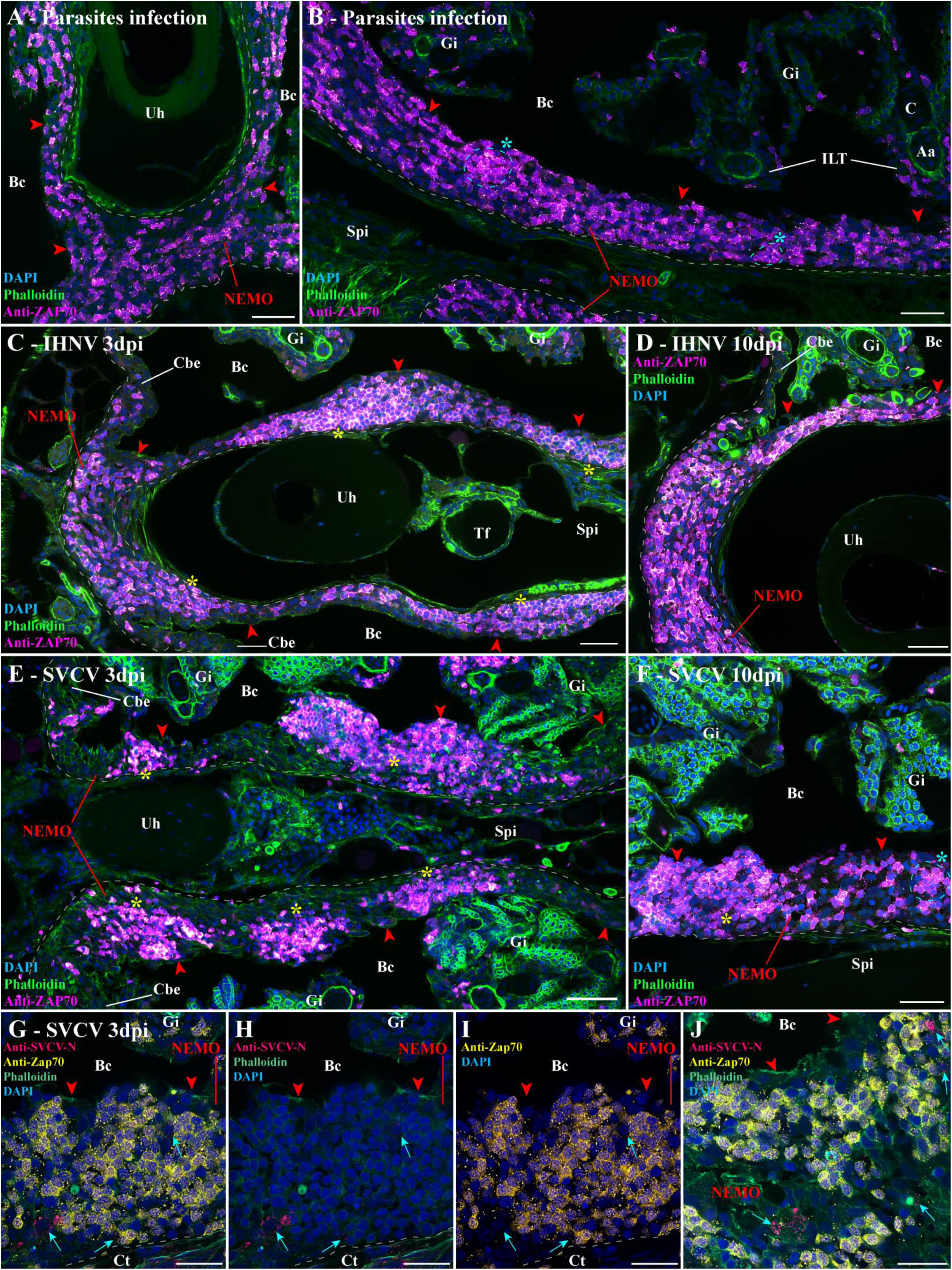
Structural response of NEMO to viral and parasitic infections. (A,B) Cryosections displaying NEMO (red arrowheads) of adult zebrafish naturally co-infected with three parasites diseases (*Pseudoloma neurophilia*, *Pseudocapillaria tomentosa*, and *Myxidium streisingeri*) labeled with anti-ZAP70 antibody (magenta hot). The distribution of ZAP70-positive cells in NEMO appears more heterogeneous than in uninfected fish. In addition, some the labeled cells formed small clusters (cyan stars). (C,D) Cryosections from adult zebrafish bath-infected for 24h with IHNV. After more prominent aggregation of ZAP70-positive cells at 3 dpi (yellow stars) (C), the distribution of T/NK cells reverted to a more homogeneous state by 10 dpi (D). (E,F) Cryosections from adult zebrafish bath-infected for 24h with SVCV. NEMO displayed striking aggregation of T/NK cells into distinct clusters at 3 dpi (yellow stars) (E). A week later, NEMO displayed both large (yellow star) and small clusters (cyan star) of ZAP70-positive cells (F). (G-J) Cryosections from zebrafish three day after SVCV infection co-labeled with anti-ZAP70 antibody (yellow) and anti-SVCV-N antibody (cherry) displaying cells loaded with virus material (cyan arrows) neighboring large clusters of ZAP70-positive cells. Annotations: Aa, Afferent artery; Bc, Branchial cavity; C, Cartilage; Cbe, Cavobranchial epithelium; Ct, Connective tissue; Dpi, day post-infection; Gi, Gills; IHNV, Infectious hematopoietic necrosis virus; ILT, Interbranchial lymphoid tissue; Spi, Sub-pharyngeal isthmus; SVCV, Spring viremia of carp virus; Tf, Thyroid follicle and Uh, Urohyal bone. Scale bars: 50 μm (E), 30 μm (A-D,F), and 20 μm (G-J).

We next studied the effect of controlled bath-infection on NEMO using two well established and commercially relevant fish pathogenic rhabdoviruses that infect tissues of the branchial cavity: Infectious Hematopoietic Necrosis Virus (IHNV) (*80, 81*) and Spring Viraemia of Carp Virus (SVCV) (*82, 83*). IHNV infection showed mild structural changes in NEMO at 3 dpi (**Fig.5 E** – yellow stars) that were evident as a seemingly deeper aggregation of ZAP70-positive cells into large clusters (yellow stars). A week later (10 dpi), these aggregates of cells had reverted toward the usual distribution of ZAP70-positive cells observed in uninfected fish (**Fig.5 F**). The effect of SVCV infection were more severe with the striking reorganization of ZAP70-positive cells into distinct large clusters at 3 dpi (**Fig.5 E, Fig.S8** – yellow stars). By 10 dpi (**Fig.5 F**), more ZAP70-positive cells, as well as smaller clusters of labelled cells (cyan star), were observed between the remaining large clusters (yellow star). When we labelled sections from the 3 dpi SVCV-infected fish with an antibody against the N protein of the virus, we detected labelled cells on the periphery of the large T/NK cells clusters (**Fig.5 G-J**). This data confirmed that the fish were successfully infected by the virus. Whether or not these labelled cells represent primarily infected cells or antigen-presenting cells that have taken-up viral material remains to be established. Collectively, these data show the involvement of NEMO in the organism response to viral pathogen infecting tissues of the branchial cavity.

In agreement with our main hypothesis, these results provide a strong evidence for the involvement of NEMO in immune responses to local infections as well as infections at distant body sites. However, while our study provides a solid foundation to study NEMO’s involvement during infection, further research is required to strengthen our understanding of NEMO’s contribution to the teleost immune response.

### NEMO, the ILTs, and ALTs as a cohesive unit of a vast lymphoid network inside the branchial cavity

Our next objective was to further investigate NEMO in the context of the branchial cavity. NEMO was intimately connected to the eight ILTs and sixteen ALTs (**Fig.1, S4**) in a way that was very striking in the 3D reconstructions (**Video.S2**). Importantly, the analysis of the anti cytokeratin staining showed that NEMO and gill lymphoid aggregates share the same network of reticulated epithelial cells (**Fig.S5 A** –red stars, **Video.S7**), indicating that NEMO, the ALTs, and the ILTs a form cohesive unit within the branchial cavity (**Fig.1**). In order to better appreciate the relation between NEMO and the lymphoid aggregates, we also looked at the structural response of the ILTs to the different infections. In parasite-infected fish, both ILTs and NEMO displayed a similar structural response as described above (**Fig.S9 A**). In both SVCV-infected and IHNV infected-fish, however, the ILTs were strongly diminished while NEMO persisted ((*36*) and **Fig.S9 B,C**) at 3 dpi, suggesting that NEMO and gills lymphoid aggregates may play a different role in cellular responses to viral infections. While NEMO is likely involved these responses based on its size, composition and location, further studies are needed to understand its precise role, as well as its relationships with ILTs and ALTs.

In addition to the ILT and ALTs, NEMO was also in continuity with regions of the cavo branchial epithelium that also contained numerous T/NK cells (**Fig.1 B** – cyan arrowheads). Using 3D multi-field of view imaging to investigate larger regions of the branchial cavity, we found that much of the cavo-branchial epithelium (**Fig.6 B,C** – cyan arrowheads) displayed a high concentration of ZAP70 positive cells, forming a vast lymphoid network within the branchial cavity that links NEMO, the sixteen ALTs, the eight ILTs and the two thymus lobes. Further analysis showed this lymphoid network extended beyond the branchial cavity region (**Fig.S10**), reaching the skin via the operculum opening (not shown), as well as the pharyngeal epithelium and the oesophagus epithelium (**Fig.S10 A**). This high T/NK cell concentration continuity further extended along the anterior segment of the pharynx and the mouth (**Fig.S10 B,C**), from which it connected with the SALT via the non-keratinized sides of the mouth opening (**Fig.S10 D**). In line with a previous study describing teleost fish SALT (*34*), it then connected with the NALT via a skin network of T/NK cells located in the basal layers of the epidermis and surrounding club cells (**Fig.S10 E-F** and **Video.S8**). Noteworthy, localized clusters of ZAP70-positive cells were observed in the epidermis of the zebrafish head, which may represent localized structured units of the SALT (**Fig.S10 C-D,F** – green stars).

**Figure 6.**
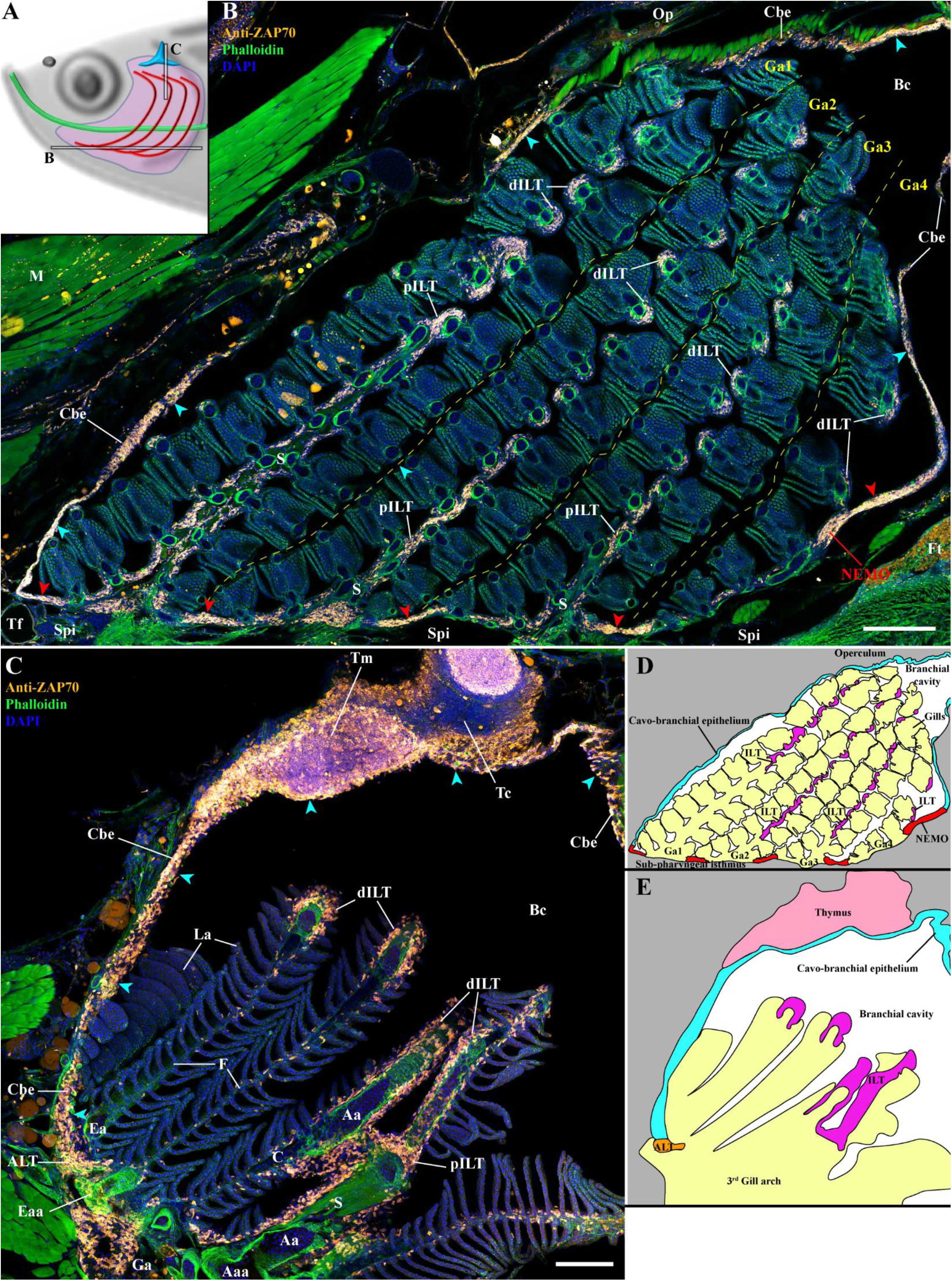
NEMO as part of a larger lymphoid network. (A) Scheme illustrating the localization the adult zebrafish NEMO images acquired from 30 μm whole-head cryosections labeled with anti-ZAP70 antibody (orange hot) to reveal T/NK cells (B-C). NEMO (red arrowheads) is part of lymphoid tissue continuity comprising a ZAP70-positive cells-rich cavobranchial epithelium (cyan arrowheads) and which connects all the lymphoid structures of the branchial cavity. (D) Scheme describing the different anatomic region displayed by the coronal cryosection presented in (B). (E) Scheme describing the different anatomic regions displayed by the transversal cryosection presented in (C). Annotations: Aa, Afferent artery; Aaa, Afferent arch artery; ALT, Amphibranchial lymphoid tissue; Bc, Branchial cavity; C, Cartilage; Cbe, Cavobranchial epithelium; dILT, distal Interbranchial lymphoid tissue; Ea, Efferent artery; Eaa, Efferent arch artery; Ft, Fat tissue; Ga, Gill arch; ILT, Interbranchial lymphoid tissue; La, Lamellae; M, Muscles; Op, Operculum; pILT, proximal Interbranchial lymphoid tissue; S, Septum; Spi, Sub-pharyngeal isthmus; Tc, Thymus cortex; Tf, Thyroid follicle and Tm, Thymus medulla. Scale bars: 200 μm (A) and 100 μm (B).

### NEMO and its cohesion with ILTs and ALTs in other teleost fish species

Our next objective was then to determine if NEMO and the lymphoid organization of the branchial cavity we described exist in other fish species. Since the zebrafish is a small representative of the cyprinid fish family, we first asked whether a larger cyprinid would share the same branchial cavity lymphoid organization. For this, we labeled wild crucian carps (*Carassius carassius*) with the anti-ZAP70 antibody, which revealed a lymphoid organization that was strikingly similar to the zebrafish (**Fig.7 A-C**). Crucian carp NEMO formed an important mass of T/NK cells wrapped around the urohyal bone that extended along the sub pharyngeal isthmus. Noteworthy, ZAP70-positive cells were also observed within the marrow of the crucian carp urohyal bone. The close association of NEMO with the ILTs and the ALTs was particularly striking, suggesting it likely remain unaffected by large body-size variations. Finally, NEMO was also connected to the ZAP70-positive cells-rich mucosal epithelium lining the branchial cavity (**Fig.7 A,B**). Intriguingly, whereas in the zebrafish NEMO was not infiltrated by endothelial vessels, NEMO of crucian carp contained clear endothelial structures in which red blood cells could be observed (**Fig.7 B,C**).

**Figure 7.**
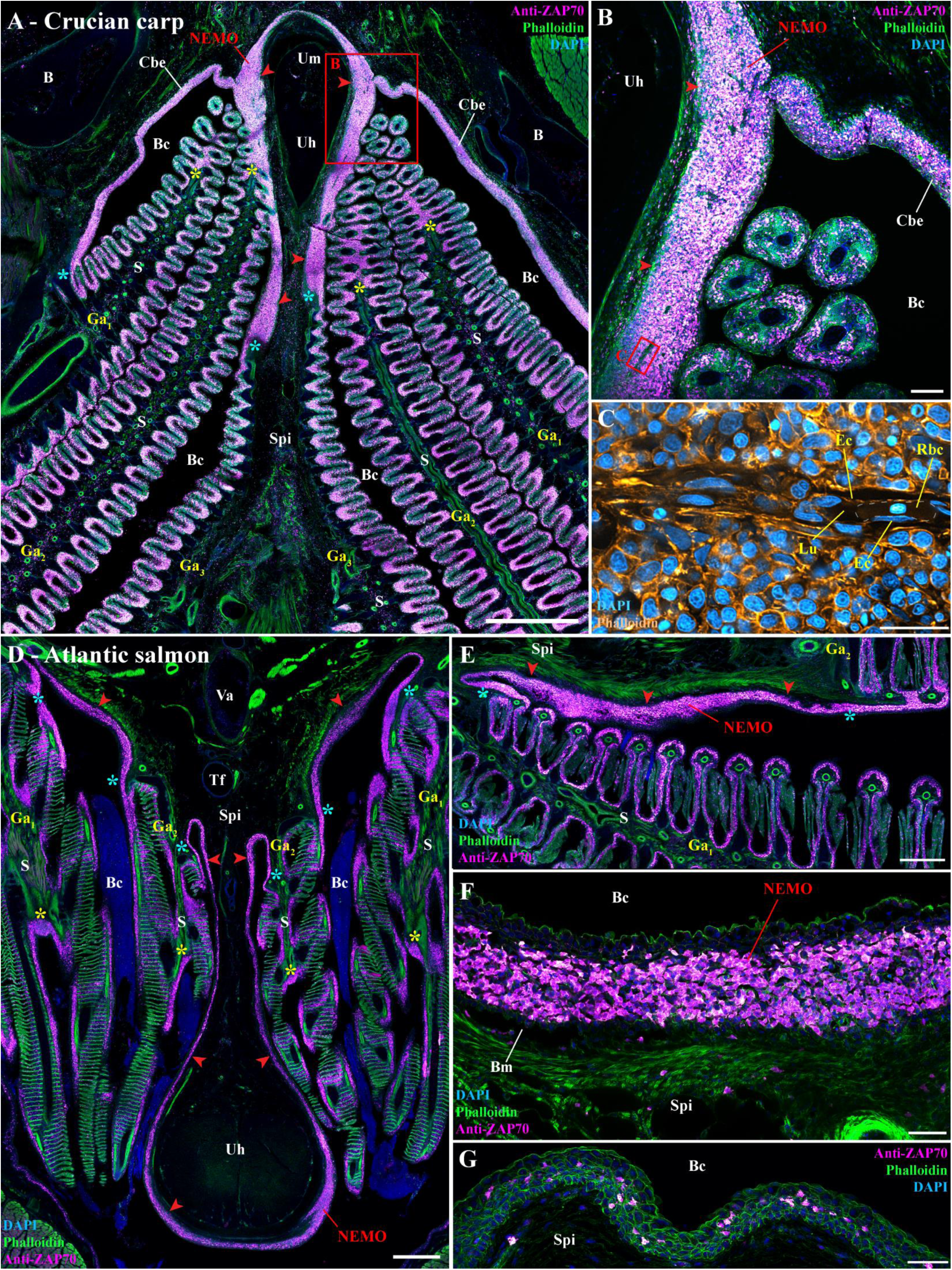
NEMO and its cohesion with ILTs and ALTs in other teleost fish species. (A-B) Cryosections from wild crucian carp labeled with anti-ZAP70 antibody (magenta hot). This larger representative of the cyprinids than zebrafish also NEMO (red arrowheads), ILTs (yellow stars) and ALTs (cyan stars), which are all interconnected by a clear lymphoid network. (C) Zoom from panel (B) in which phalloidin (orange hot) and DAPI (cyan hot) staining revealed the presence of endothelial vessels containing red blood cells. Transversal (D) and coronal (E) cryosections of an adult Atlantic salmon (1.5 kg) labeled with anti-ZAP70 (magenta hot) displaying NEMO (red arrowheads), ILTs (yellow stars), ALTs (cyan stars). As for cyprinids, NEMO (red arrowheads) of this representative of the salmonid family is located at the ventral convergence of gill arches, interconnected with gills lymphoid aggregates, and closely associated with the urohyal bone. It is particularly enriched in ZAP70-positive cells (F). In contrast, the mucosal epithelium lining the region directly anterior to the urohyal bone only displayed scarce T/NK cell (G). Annotations: B, Bone; Bc, Branchial cavity; Bm, Basement membrane; Cbe. Cavobranchial epithelium; Ec, Endothelial cell; Ga, Gill arch; Lu, Lumen; Rbc, Red blood cell; S, Septum; Spi, Sub-pharyngeal isthmus; Tf, Thyroid follicle; Uh, Urohyal bone; Um, Urohyal marrow and Va, Ventral aorta. Scale bars: 1000 μm (A), 500 μm (D), 400 μm (E), 100 μm (B), 50 μm (F,G), and 20 μm (C).

We then extended our investigation to the adult Atlantic salmon (*Salmo salar*), a representative salmonid (**Fig.7 D-G**). While adult Atlantic salmon is anatomically divergent from zebrafish and crucian carp (E.g. open branchiostegal rays), and displays different adaptations (E.g. Atlantic salmon is an anadromous marine fish), we found a structure very similar to cyprinid NEMO in that it is also wrapped around the urohyal bone and intimately connected to the gill lymphoid aggregates (**Fig.7 D,E**). In contrast to the abundance of T/NK cells in Atlantic salmon NEMO (**Fig.7 F**), fewer ZAP70-positive cells were observed in the mucosal epithelium anterior to the urohyal bone (**Fig.7 G**). During our literature search, we came across one amazing old research article from 1939 on the Atlantic salmon thyroid in which we could recognize a structure that was highly reminiscent of NEMO (*84*). In that paper, the putative NEMO can be observed at younger stages of Atlantic salmon development such as fry and parr. We could also recognize a similar structure reminiscent of NEMO within the second figure of a study on the thyroid of three-spined stickleback (*Gasterosteus aculeatus*), a representative of the gasterosteid (*85*), a family belonging to the most derived clade of euteleost, the percomorphs.

Collectively, this probing comparative study indicates that NEMO and the branchial cavity lymphoid organization we describe here in zebrafish are conserved among members of the cyprinid and salmonid families, the two most extensively studied and farmed families worldwide, as well as in some percomorphs. Moreover, our data suggest that cohesiveness with gill lymphoid aggregates and the anatomical associations with the urohyal bone and sub pharyngeal isthmus might represent typical features of NEMO across different teleost species.

## DISCUSSION

In the present study, we investigated the structural organization of lymphoid tissues in the zebrafish branchial cavity. We show here that the tissues lining the branchial cavity, which are highly exposed to the pathogens of the aquatic environment, present a complex network of connected immune tissues. Advanced imaging of this region led us to identify a novel mucosal lymphoid organ, which we tentatively named the Nemausean Lymphoid Organ (NEMO), that is associated to the fish pharyngo-respiratory tract. Detailed investigations of its structural organization by light and electron-microscopy provided a solid foundation to characterize NEMO as a lymphoid organ, such as the presence of an intricate network of reticulated epithelial cells. Importantly, NEMO was a prominent constituent of a cohesive unit, formed with ILTs and ALTs, of a lymphoid network interconnecting all the lymphoid structures in the branchial cavity. This highlights a new level of integration of the surveillance and defense system associated to the fish pharyngo-respiratory tract. Given the central localization of NEMO within the branchial cavity, we hypothesized that NEMO could constitute a secondary lymphoid organ in which immune responses would occur. This idea was supported by the characterization of its immune cell populations that showed a high-enrichment in T/NK cells and B cells mixed with antigen-presenting cells, and by the lack of expression of the recombinase required for V(D)J rearrangements, RAG2. NEMO also contains ZAP70^+^ PCNA^+^ proliferating T cells, and ZAP70^+^ MHCII^+^ and ZAP70^+^ TNFα^+^ activated T cells, further strengthening the hypothesis that it is a secondary lymphoid organ. Following infection by different pathogens, NEMO underwent structural changes involving the formation of ZAP70^+^ cells clusters, suggesting direct involvement in immune responses.

During our investigations, we could make an initial appreciation of NEMO’s ontology using our data and the zebrafish histology atlas (https://bio-atlas.psu.edu/zf/progress.php), and discovered that NEMO appears surprisingly late during development. Whereas the main lymphoid organs (thymus, spleen, kidney) are already present by the two first weeks of the zebrafish development (*86, 87*), NEMO likely appears around the 4^th^-6^th^ weeks of development, right after the larval-juvenile transition stage. Thus, NEMO’s development coincides with the emergence of a fully mature adaptive immunity (*87, 88*). The gill lymphoid aggregates (ALT and ILT), to which NEMO is tightly connected, also appear during the same time window, or possibly just after the development of NEMO. The appearance of these lymphoid structures is particularly striking when one considers that in 3 wpf zebrafish, besides for the thymus lobes, the branchial cavity contains very few T/NK cells. Further work will be required to understand the developmental mechanisms involved in this drastic transformation of the immune landscape of the branchial cavity, and its impact on the fish susceptibility to infections. Interestingly, it is well known that fish fry are highly sensitive to many viral and bacterial infections compared to adults. In mammals, thyroid hormones have a significant impact on lymphoid organs’ development and biology (*89, 90*). For example, early removal of the thyroid in rats impairs lymphoid organs morphology and functions (*91*). In zebrafish, thyroid hormones are important for the larval-juvenile transition stage, and they also influence the size of the thymus (*92–94*). The possibility that thyroid hormones may impact NEMO’s development and, more generally the relocation of the T-cells in the branchial cavity, would deserve some attention. The presence of a recently reported zebrafish cell population sharing molecular markers with mammalian lymphoid tissue inducer cells (LTi) (*15*), a type of innate lymphoid cell that is involved in the formation of certain secondary lymphoid organs in mammals (*95*), should also be investigated in NEMO and in the branchial cavity at the end of the larval-juvenile transition stage.

An interesting question is whether NEMO has counterparts among vertebrates or if this organ represents an independent solution developed by certain fish species to combat infectious diseases. The evolutionary history of lymphoid organs has been investigated and intensely debated using various terminologies and classification criteria (*3–7, 9, 10*). In addition to the thymus, the presence of lymphoid aggregates associated to the pharyngeal region has been reported in mammals, birds, reptilians, and amphibians, but so far not in teleostean or chondrostean fish. Over a century ago, Salkind mentioned the existence of lymphoid concentrations within the lower pharynx of Brook lamprey (*Lampetra planeri*), an agnathan; he considered that these structures were distinct from the presumptive thymus (*9*). The absence of prior recognition of a prominent pharyngeal lymphoid structure such as NEMO underscores that teleost fish lymphoid organization remains imperfectly comprehended. Collectively, a fascinating question that arises from this work is whether the formation of mucosal secondary lymphoid structures within the pharyngo-respiratory region is a conserved developmental program shared by vertebrates.

In mammals, the nasopharyngeal area is protected by a set of mucosal secondary lymphoid organs called tonsils, which collectively form the Waldeyer’s tonsillar ring (*96*). Several lines of evidence support the hypothesis that NEMO may represent a fish analogue of mammalian tonsils. NEMO, like the tonsils, is a mucosal lymphoid organ located within the pharyngo respiratory tract, which is exposed to the external environment. As tonsils, NEMO constitutes a mass of lymphoid cells structured by an intricate network of reticulated epithelial cells. Moreover, in certain fish species NEMO exhibits clear signs of vascularization. NEMO is part of a larger lymphoid network containing 25 lymphoid structures (8 ILTs, 16 ALTs, and NEMO) that are strategically positioned within the pharyngo-respiratory region at sites exposed to antigens encountered during feeding and breathing. This arrangement might represent in fish a distant functional analogue of the Waldeyer’s ring of tonsils. Finally, as palatine tonsils, NEMO appears relatively late in development at a site associated with the 2^nd^ pharyngeal pouch (*97– 100*). However, mammalian tonsils display a more complex structural organization than NEMO, such as well-defined T cell and B cell zones. Similarly, the presence of germinal-like centers in NEMO remains to be explored.

The analysis of NEMO cellular composition is particularly significant for our understanding of teleost fish immunology. We showed that NEMO contains more T/NK cells than the spleen, which is considered as the primordial secondary lymphoid organ in fish (*4*), suggesting it may play an essential role for the homeostasis of adaptive immunity. Ablation of NEMO or T-cell depletion experiments, for example using rag2 mutant zebrafish (*101*), could provide valuable information on NEMO’s function. In addition, we showed that NEMO also possess a significant plasma/B cells population. Secretory antibodies plays an important role in the maintenance of the branchial cavity homeostasis, as shown by Xu *et al* (*102*), the experimental depletion of secretory IgT immunoglobulin in rainbow trout induced gill dysbiosis, inflammation and tissue damages. Further studies would have to clarify the role of NEMO in immune system regulation and the screening of external pathogens from local microbiota.

The infection experiments we performed have been key to propose NEMO as a secondary lymphoid organ. After 3 day of infection by SVCV, NEMO displayed a rearrangement of T/NK cells into large clusters surrounded by cells carrying virus material. Further studies would have to determine the nature of these concentrations of lymphoid cells we observed in NEMO and if they represent structures favoring processes of adaptive immunity such as antigen-presentation. It would be particularly interesting to determine if they are associated to the formation of melanomacrophage center (MMC), which are immune structure that have been suggested as potential fish analogue of mammalian germinal center (*103*).

Our observations revealed a structural cohesion between NEMO, ALT, and ILTs, all sharing the same network of reticulated epithelial cells. The ILT was the first structured mucosal lymphoid tissue discovered in fish. Studies in salmonids showed that it may represent a non conventional secondary lymphoid tissue (*26, 73, 104, 105*). This was followed by our description of the ALT in zebrafish. Consistent with studies in salmon, the zebrafish ILT and ALT displayed features of secondary lymphoid organ. In this study we showed that ILTs and ALTs are not just a set of 24 distinct lymphoid structures, they are bound together by a more prominent lymphoid organ that also possess features of secondary lymphoid organs, NEMO. Intriguingly, NEMO, the ALTs and ILTs did not display a similar structural response to IHNV and SVCV infections in adult zebrafish (*36*). The zebrafish ILT and ALT are first strongly reduced by 3 dpi, which is consistent with a reduction of the ILT described in Atlantic salmon infected with infectious salmon anemia virus (*73*), whereas NEMO persisted. This constitutes a major difference between NEMO and the gill lymphoid aggregates, which indicate they likely bear different immune functions. However, NEMO and the ILT displayed a similar rearrangement of ZAP70 cells in the fish naturally co-infected with multiple parasites. Further studies would have to determine if the gill lymphoid aggregates are constituents of NEMO.

Our investigations of the whole branchial cavity revealed the presence of a vast lymphoid network that links NEMO, ILTs, ALTs and thymus lobes altogether into a sophisticated super structure of the fish immune system, which suggest the branchial cavity may act as a lymphoid nexus. The implications of this lymphoid super-structure for the homeostasis of the fish immune system and its interactions with other tissues/organ remains to be deciphered.

In this study, we recognized NEMO in representatives of distant teleost fish species families: two cyprinids, one salmonid, and a Percomorph. This observation suggests that NEMO is likely present in most fish families. We anticipate a significant level of structural variability in NEMO across teleost fish species, given the taxonomic and morphologic diversity within this group. However, our data suggest that its position around the urohyal bone and at the convergence of gill arches along the sub-pharyngeal isthmus are conserved features. Importantly, as in zebrafish, the integral unit formed by NEMO and the gill lymphoid aggregates was clear in all the analyzed fish species, suggesting that the NEMO/ALT/ILT apparent unity might be an essential arrangement for immune responses in the branchial cavity. Taking into account our previous comparative studies, ILTs seems only present within the gills of “basal” teleost, whereas ALT and NEMO could be observed in all the analyzed teleost fish species so far. It would be of particular interest to investigate if the morphology of the branchial cavity, and more specifically the evolution of the branchiostegal rays (*49*), influence the existence or the morphology of NEMO and its unity with gill lymphoid aggregates.

## CONCLUSION

Altogether, our study provides new insights about the teleost fish immune system and its structural organization. We identified a novel lymphoid organ within the pharyngo-respiratory region of adult zebrafish and other teleost species, which we named “Nemausean Lymphoid Organ (NEMO)”. Our investigations led to the idea that NEMO is a fish mucosal secondary lymphoid organ that shows features of mammalian tonsils. Intimately associated with gill lymphoid aggregates, NEMO appears as a potential key lymphoid hub coordinating lymphocyte traffic and defense mechanisms within the fish respiratory mucosa. Collectively, our findings contribute to a better understanding of the evolution of the vertebrate immune system, and provide new insights in fish immunology. Gill immunity is of growing importance both for future vaccines in aquaculture and for the development of disease models in zebrafish.

## MATERIALS AND METHODS

### Animal Care and Ethic Statement

Experiments were conducted in compliance with the animal care guidelines, ethical standards and legislation of the European Union, France and Norway, and in consultation with local ethics committees. Animal experiments performed in the present study were carried out at the IERP fish facilities (building agreement n°C78-720, doi.org/10.15454/1.5572427140471238E12) of the INRAE Research Center at Jouy-en-Josas, France, in compliance with the recommendations of Directive 2010-63-EU on the protection of animals used for scientific purposes. These infection protocols were authorized by the institutional review ethics committee, COMETHEA, of the INRAE Research Center. Authorizations were approved by the French Ministry of Agriculture (authorization number APAFIS# 22150-2019062811528052).

Animal experimentations, handling, and euthanasia were performed by well-trained and authorized staff. Specimen were euthanized using an anesthetic overdose of buffered tricaine.

The experiments were performed using AB wild-type zebrafish (around 1 year, unless specified) (N=36) and the following transgenic lines: *Tg(lck:EGFP)* (N=3) (*16*)*, Tg(fli1a:EGFP)^y1^* (N=3) (*57*), *Tg(mpx:GFP)^i114^* (N=3) (*63*), *Tg(mhc2dab:GFP)^sd6^* (N=3) (*64*), Tg(mfap4:mCherry-F)^ump6^ (N=6) (65), Tg(Cau.Ighv-ighm:EGFP)^sd19^ (N=3) (71), Tg(rag2:DsRED) (N=3) (72), Tg(cd41:GFP) (N=3) (74), and Tg(tnfα:eGFP-F)^ump5^ (N=3) (78).

The study includes three laboratory grown adult zebrafish (1 year) naturally co-infected with *Pseudoloma neurophilia*, *Pseudocapillaria tomentosa*, and *Myxidium streisingeri*, provided by the Oehlers’ laboratory (SINGAPORE). The fishes were held at the IMCB Zebrafish facility under IACUC approval 211667, and were sampled as part of a culling following veterinarian diagnosis. Animal handling and euthanasia were performed in accordance with Singaporean regulations.

Two healthy adult Atlantic salmon (weight: 1500g), laboratory-raised by NIVA (Solbergstrand NORWAY), were provided by PHARMAQ, a division of Zoetis. The fish were handled and euthanized in strict accordance with Norwegian legislation by authorized staff.

Three wild crucian carps (40g: both sex), captured using nylon net in October 2020 in Tjernsrud pond (Oslo-NORWAY), with a healthy appearance upon sampling were provided by the Lefevre-Nilsson group from the University of Oslo. Specimen were sampled and euthanized in compliance with Norwegian animal welfare laws (Dyrevelferdsloven), carried out as part of the authorized project FOTS permit ID 16063, and following the instruction about use of animal for research (Forskriften om bruk av dyr I forsøk).

### Infection experiments with SVCV and IHNV

Spring Viremia of Carp Virus (SVCV) and Infectious Hematopoietic Necrosis Virus (IHNV) infectious challenges were carried out on wild-type zebrafish of the AB strain, aged 16 months and weighing 0.8g (+/-0.03g). Fish were acclimatized for 48h at 22°C (pH 7, conductivity 200µS/cm²) in 1.5L aquaria. Two groups of eight fish each were then infected by immersion for 48 h using the reference SVCV strain VR-1390 (PMID: 29114248) and the IHNV strain 25-70 adapted to 25°C (*108*) and the IHNV strain 25-70 adapted to 25°C (*109*), at a final concentration of 104 PFU/ml. The water flow was stopped for 48 h, followed by daily water change. Fish were euthanized and sampled at 3 and 10 days post infection by IHNV, and at 3 and 10 dpi by SVCV. Non-infected controls were prepared in parallel (n = 4).

### Electron microscopy

Juvenile zebrafish (9 wpf, N=3) were euthanized and immediately immersed in 20mL of fixative (4% Formaldehyde, 0.8% Glutaraldehyde GA, in 1X PHEM buffer (*110, 111*) pH7.2 in fish water) for 24h at room temperature (RT), followed by a 24h incubation at 4°C. Samples were quenched in 100 mM glycine for 2h at RT and rinsed with 100 mM sodium bicarbonate buffer (pH 6.5). For postfixation, samples were incubated on ice with a solution of 2% osmium tetroxide and 1.5% potassium ferricyanide in 100 mM sodium bicarbonate buffer (*112*), rinsed 5 times with 100mM sodium bicarbonate buffer and 2 times with 50 mM maleate buffer (pH 5.15), and incubated in 2% uranyl acetate in 50 mM maleate buffer for 3h (*113*). Following washes in 50 mM maleate buffer, gradual dehydration was achieved by “progressive lowering of temperature” (*114*) using the following sequential incubations: 1h 30% ethanol on ice, 1h 50% ethanol on ice, 30min 1% uranyl acetate in 70% ethanol at -20°C, 1h 70% ethanol at - 20°C, 90min 80% ethanol at -30°C, 90min 90% ethanol at -30°C, 2h 96% ethanol at -30°C, 16h 100% ethanol at -30°C, 3 times 2h 100% dry ethanol (3Å molecular sieve (*115*)) at -30°C, 2 times 30min dry acetone at -30°C, and 14h 25% EPON in dry acetone at RT. EPON (*116*) was prepared with a ratio of 3:7 (DDSA:NMA) containing 1% DMP-30. Specimen infiltration was done by a 24h incubation in 100% EPON at RT. Samples were then embedded in fresh EPON using flat embedding molds and oriented after 3h of polymerization at 60°C. Finally, polymerization was performed for 48h at 60°C followed by a 24h curing period at RT. Targeted trimming was aided by staining semithin (300 nm) sections with 0,1% toluidine blue in borate buffer (pH 11) at 80°C (*117, 118*) to facilitate the orientation in the sample. Samples were sectioned at 60 nm thickness on a Leica UCT ultramicrotome using Diatome 45° ultra knifes and mounted on carbon coated, formvar film on 2 mm single hole copper grids. Sections were then stained with 4% uranyl acetate in 50% methanol for 1h (*119*), followed by a 20 seconds incubation with Reynolds lead citrate (*120*). Images were acquired at 120 kV with a Jeol JEM-1400 electron microscope using a Tvips 216 camera. The manually recorded images were aligned using the plugin big stitcher in ImageJ, montaged using gimp for layer projection, and colored using photoshop CS6. If applicable, generation of a tile pyramid and visualization via java was done using Open-Seadragon and Openlayers on a basic html site.

Several ultrastructure maps of NEMO are available at the following internet addresses: (https://wohlmann.github.io/2019019_004_M1c/) (https://wohlmann.github.io/2019019_004_N2/)

### 3D reconstruction of the zebrafish NEMO

Following euthanasia, adult zebrafish heads (15 wpf, N=3) were fixed in a solution of 4% methanol-free formaldehyde (Thermofisher) in 60 mM HEPES buffer (pH 7.4) for 24 h at room temperature, followed by a 3 days incubation at 4°C. For decalcification, samples were incubated in a solution of 13% EDTA (pH 7.8), 0,1% tween (Sigma Aldrich) and 1% triton X-100 (Sigma Aldrich), in ddH_2_O for 5 days at RT under gentle rocking. Samples were then saturated for 24h at RT in a blockaid solution (Thermofisher) with 0,5% triton X-100 and 0,1% tween. T/NK lymphocyte were labeled using a rabbit anti-ZAP70 monoclonal antibody (99F2 – Cell Signaling) diluted at 1:600 in Pierce™ Immunostain Enhancer solution (Thermofisher) complemented with 0,5% triton X-100 and 0,1% Tween for 5 days at RT under gentle rocking. Samples were then washed several times at RT in 1X PHEM buffer (60 mM PIPES, 25 mM HEPES, 10 mM EGTA and 2 mM MgCl2 in ddH_2_O – pH 7.4 (*110, 111*)) with 0,5% triton X-100 and 0,1% Tween (PHEM_t-tw_), and incubated with goat anti-rabbit-Alexa647 (Jackson ImmunoResearch) diluted at 1:400 in 1X PHEM_t-tw_, complemented with phalloidin-TRITC (Sigma Aldrich) at 3U/mL and DAPI (Thermofisher) at 5 µg/mL, for 5 days at RT under gentle rocking. Samples were then first rinsed with 1X PHEM_t-tw_ and then 1X PHEM. Samples were stored at 4°C in 1X PHEM until further processing. One sample was then sliced and mounted onto a coverslip with slowfade glass mounting medium (Thermofisher) to control the quality of the labeling and for wholemount imaging of the skin covering the head.

Tomography was performed using an automatized Zeiss LSM880 confocal microscope coupled with a vibratome (Microm HM 650V from Thermo Scientific). For sectioning, ZAP70-labeled zebrafish heads were embedded in 6% agarose in water into a 1 cm square plastic chamber and orientated for appropriate cross-sectioning with rostral side on top. Once set, agarose box was resized using a razor blade and was attached with superglue on a metal surface with rostral side orientated on top and placed into a tank filled with water. The following automated process was then applied: a 80 µm thick layer was removed from the surface of the block containing the samples by the vibratome, followed by the immediate imaging of the newly exposed surface with the Zeiss LSM 880 microscope. Image were acquired in confocal mode, with a 20x Plan Apo 1.0 NA water immersion objective, the wavelength 633 nm for excitation and 660-711 nm band for emission and the wavelength 561 nm (Argon Laser) and 561-630 nm band (GaAsP detector), sequential mode, mosaic of 12 x 12 fields, stack of 102 µm total volume and 6 µm steps. Imaging steps were repeated The 3D files generated from acquisitions were processed using Image J for alignment, stitching, and cropping. NEMO, the thymus lobes, the ALT, and the ventral end of the gill arches, were manually segmented on each single layer based on the phalloidin and anti-ZAP70 signals using Imaris in order to assemble the different 3-D reconstructions. IMARIS was also used to generate the 3D videos.

### Immunofluorescence - Cryosections

Following euthanasia, whole adult zebrafish and dissected lower pharyngeal areas of both Atlantic salmon and crucian carp, were fixed in a solution of 4% methanol-free formaldehyde (Thermofisher) in 60 mM HEPES buffer (pH 7.4) for 24 h at room temperature, followed by a 3 days incubation at 4°C. Atlantic Salmon and crucian carp samples were decalcified with a 5 days incubation in a solution of 13% EDTA (pH 7.8) in ddH_2_O at RT under gentle rocking.

Samples were cryoprotected by two incubations in a solution of sucrose at 32% in ddH_2_O, until the specimens sunk to the bottom of the recipient, and embedded in Tissue-Tek O.C.T. Compound (Sakura Finetek USA, Mountain View, CA, USA). Samples were flash-frozen in isopentane, and sectioned using a CM1950 cryostat (Leica, Wetzlar, Germany). The resulting 30 µm cryosections were collected on Superfrost Plus slides (Thermofischer) and stored at - 20°C. Samples used for stereology analyses were sectioned and recovered in standardized uniform random way.

Following the protocols detailed in (*36*), immunofluorescence as follow. Briefly, following saturation in blockaid solution (Thermofisher), slides were incubated with one or several of the following primary antibody / lectin: 1:300 rabbit anti-ZAP70 monoclonal antibody (99F2 – Cell Signaling), 1:40 cytokeratin Pan Type I/II mouse monoclonal antibody cocktail (Thermofisher), 1:300 mouse anti-PCNA monoclonal antibody (PC10 – Thermofisher), 1:200 mouse anti-BTK monoclonal antibody (D6T2C – Cell Signaling), and 1:20 mouse anti-SVCV-N monoclonal antibody (BIO 331 – Bio-X Diagnostics). 1:200 peanut agglutinin lectin coupled with alexa594 (Thermofisher). When necessary, sections were incubated with one or several of the following cross-adsorbed secondary antibodies at 1:250: Goat anti-rabbit-Alexa647 (Jackson ImmunoResearch), Goat anti-mouse-Alexa647 (Jackson ImmunoResearch), and Goat anti mouse-Alexa594 (Jackson ImmunoResearch). Where relevant, secondary antibodies or lectin were co-incubated with fluorescent phalloidin (TRITC or FITC labeled – Sigma Aldrich) at 3U/mL, and DAPI (Thermofisher) at 5 µg/mL. Slides were mounted with coverslips using prolong-glass mounting medium (Thermofisher).

### Imaging and Image analysis

3D images were acquired with the Zyla camera of a dragonfly 500 spinning disk confocal microscope (Andor, Belfast, UK), with 40 µm pinholes and either a 20×/0.75-dry objective or a 60×/1.4-oil-immersion objective. Acquisitions, stitches and deconvolutions (14-16 iterations) were performed using in-build features of the Fusion software. Image analysis were carried out using IMARIS and ImageJ/FIJI softwares. The acquisition and analyses of images were made at the NorMIC imaging platform (University of Oslo, NORWAY).

The average volume of a zebrafish T/NK cell was quantified using IMARIS by reconstructing the 3D structure of 15 random ZAP70-positive cells from 3D images of NEMO cryosections coming from 3 different fish.

The volume of NEMO, and spleen, occupied by ZAP70-positive cells was calculated with “point counting stereology” (*121*) using ImageJ to generate randomly placed 500 µm^2^ and 100 µm^2^ grids on single optical section images of the anterior segment of NEMO collected from 3 different fish. The amount of T/NK cells was then calculated by multiplying the total volume of the organ by the fraction of the volume occupied by ZAP70-positive cells, which was then divided by the average volume of a single T/NK cell.

Cell counting was performed manually using the “Cell counter” plugin on ImageJ on single optical section images of the anterior segment of NEMO coming from 3 different fish. Graph were generated using the software GraphPad Prism 7.

## ACKNOWLEDGEMENT

We thank the Oslo NorMIC imaging plateform (O. Bakke, F. Skjeldal and L. Haugen), the IBV Electron-Microscopy facility (N. Roos), the histology platform of the FYSCELL section (T. Klungervik), the NCMM zebrafish facility (C. Esguerra, AC. Tavara), the A*STAR IMCB Zebrafish Facility, as well as K.Zulkefli and the Ella Maru studio (E. Marushchenko and K. Zvorykina) for the scientific illustrations. We would also like to ackownledge the imaging facility BioCampus Montpellier Ressources Imagerie (MRI) (M. Dejean and M. Lartaud), member of the national infrastructure France-BioImaging supported by the French National Research Agency (ANR-10-INSB-04, “Investments for the future”). We also address a special thanks to C. Wiik-Nielsen and H. Hardersen at PHARMAQ, part of Zoetis, for providing us with salmon samples, and to L. Gerber, G. Nilsson, and S. Lefevre (University of Oslo) for providing crucian carp samples. We would also like to acknowledge A. Zapata, L. Du Pasquier,

A. Dalum, S. Khan, A. Kraus, M. Groß, E. Davydova, B. Mathiesen, A.López-Porras, JP. Levraud, G. Lieschke, H. Winther-Larsen, N. Steinel, P. Elk, R. Hodge and E. Teyssier for their highly appreciated help, discussion, and support. Finally, we thank all the persons that have been involved in the creation of the different genetically modified zebrafish strains.

We thank the Norwegian Research Council for funding (No 144642 and No 329478). In addition, PB and BV were supported by the Agence Nationale de la Recherche (France) (project ANR-21-CE35-0019), and by the ERANET project Nucnanofish. MN-C was supported by the french Agence Nationale de la Recherche [ANR-19-CE15-0005-01, MacrophageDynamics].

## AUTHOR CONTRIBUTIONS

Conceptualization: PB, GG, and JR. Investigation: MN-C, JW, DR, and JR. Methodology: MN-C, JW, DR, GG, and JR. Validation: JR. Writing - Original draft: GG, PB, and JR. Writing - Review & Editing: MN-C, JW, DR, SO, FJ, SQ, BV, GW, IS, EK, PB, GG, and JR.

Visualization: JW and JR. Supervision: MN-C, IS, PB, BV, FJ, SQ, GG, and JR. Project administration: JR. All authors contributed to the article and approved the submitted version.

## CONFICT OF INTEREST

The authors declare that the research was conducted in the absence of any commercial or financial relationships that could be construed as a potential conflict of interest.

## ABBREVIATIONS

Aa: Afferent artery
Aaa: Afferent arch artery
ALT: Amphibranchial Lymphoid Tissue
B: Bone
Bc: Branchial cavity
Bm: Basement membrane
BTK: Bruton Tyrosine Kinase
Bv: Blood vessel
C: Cartilage
Cbe: Cavo-branchial epithelium
Ct: Connective tissue
Cvs: Central venous sinus
d-ILT: distal Interbranchial Lymphoid Tissue
Dpi: Day post-infection
Ea: Efferent artery
Eaa: Efferent arch artery
Eav: Endothelial anastomotic vessels
Ec: Endothelial cell
EM: Electron Microscopy
Ft: Fat tissue
Ga: Gill arch
GALT: Gut-Associated Lymphoid Tissue
Gi: Gills
GIALT: Gill-Associated Lymphoid Tissue
H&E: Hematoxylin and Eosin
HK: Head-Kidney
ILT: Interbranchial Lymphoid Tissue
IHNV: Infectious Hematopoietic Necrosis Virus
La: Lamellae
Lu: Lumen
M: Muscles
MALT: Mucosa-Associated Lymphoid Tissue
NALT: Nasal-Associated Lymphoid Tissue
NEMO: Nemausean lymphoid Organ
NK: cell Natural Killer cell
Op: Operculum
PCNA: Proliferating Cell Nuclear Antigen
p-ILT: proximal Interbranchial Lymphoid Tissue
Rbc: Red blood cell
S: Septum
SALT: Skin Associated Lymphoid Tissue
Sk: Skin
Spi: Sub-pharyngeal isthmus
SVCV: Spring Viremia of Carp Virus
Tc: Thymus cortex
Td: tendon
TEM: Transmission Electron Microscopy
Tf: Thyroid follicle
Tm: Thymus medulla
Tu: Tubule
Uh: Urohyal bone
Um: Urohyal marrow
Va: Ventral aorta
Wpf: week post-fertilization
ZAP70: Zeta-chain-Associated Protein kinase 70.

## SUPPLEMENTARY FIGURES AND VIDEO LEGENDS

**Figure S1.**
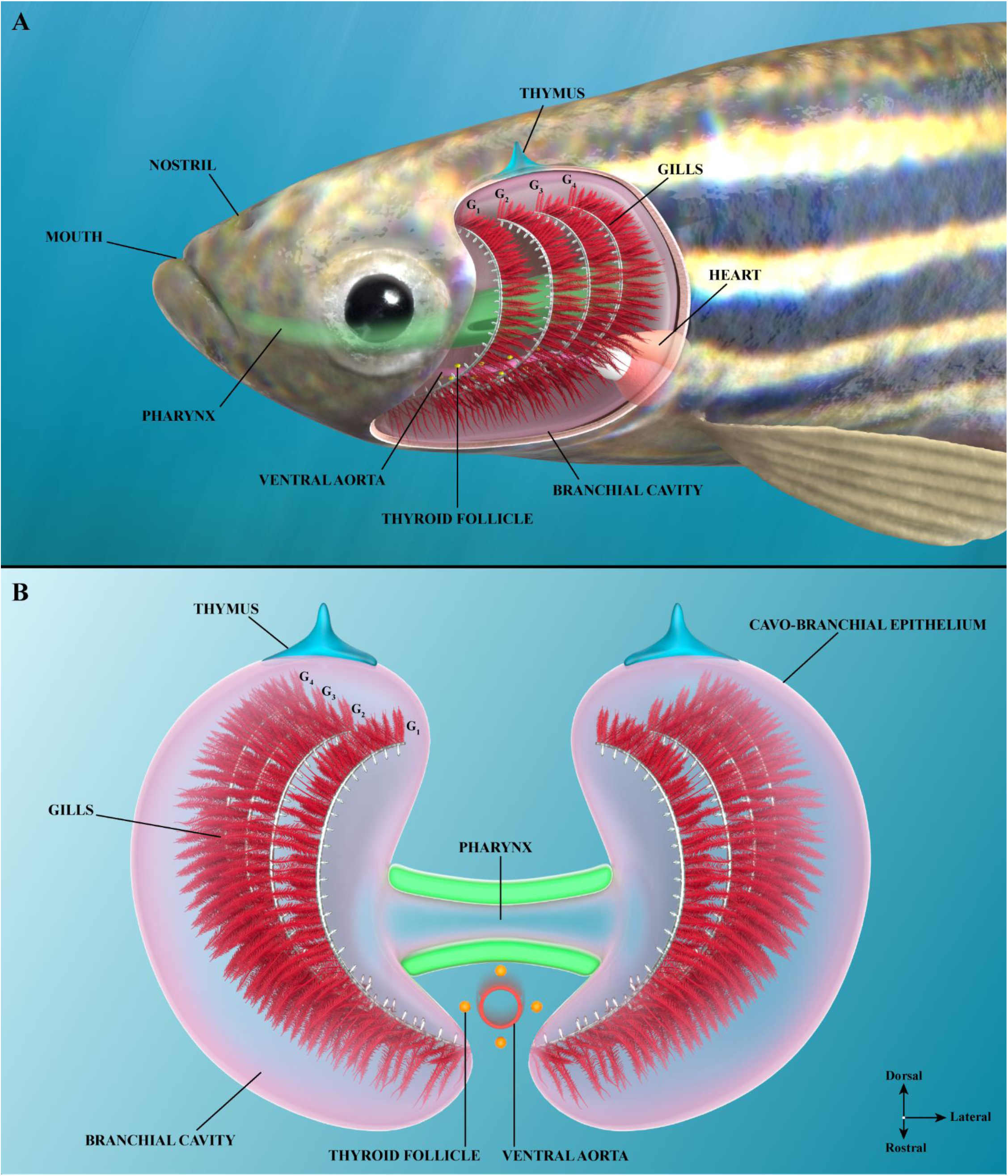
Organization of the adult zebrafish branchial cavity. Illustrations of the branchial cavity tissue organization as seen from the side (A) or from a front view (B). G1-4: First to fourth gill arch. Illustrations made by Ella Maru studio.

**Figure S2.**
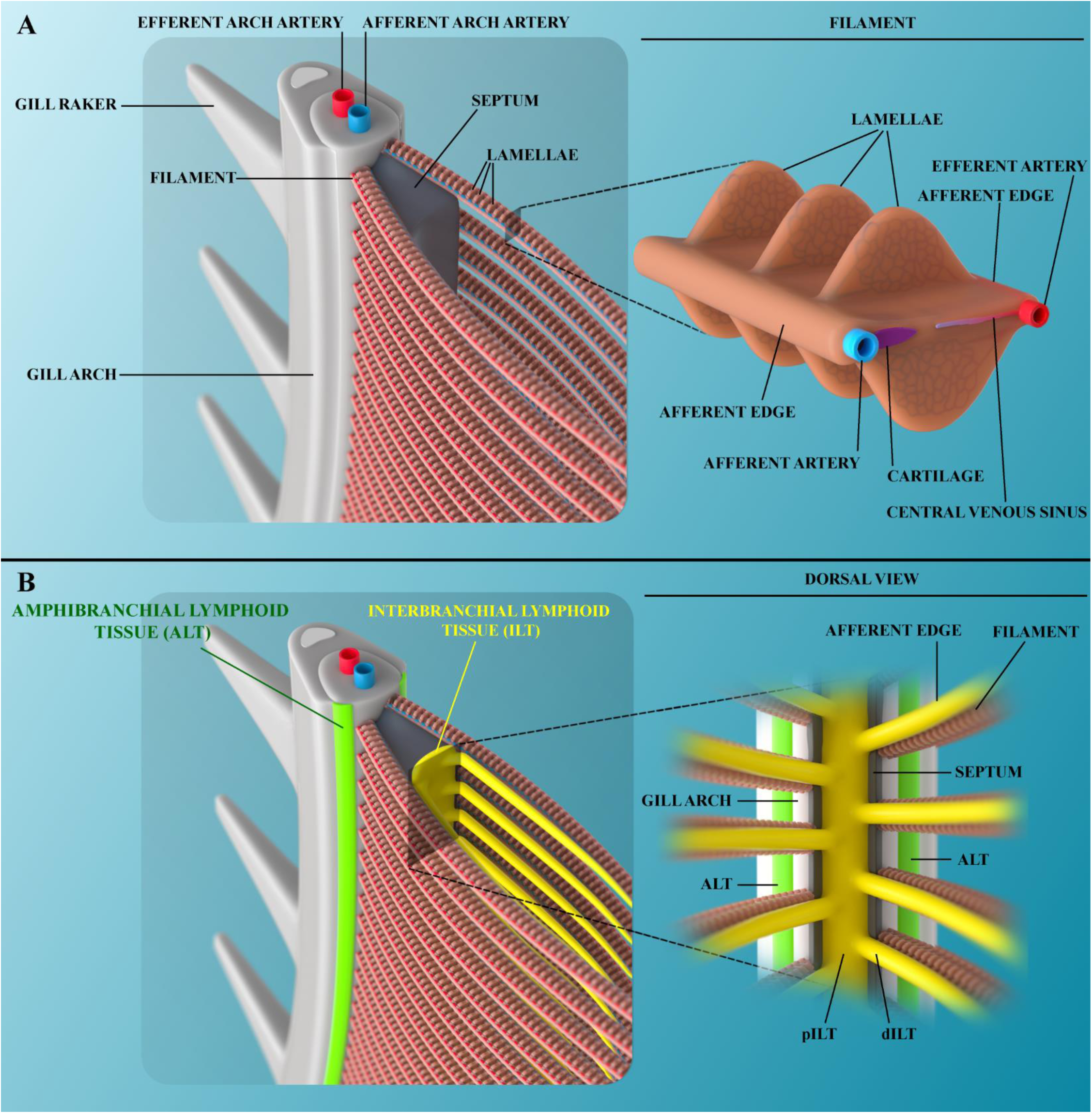
Organization of the adult zebrafish gills. Illustration of a gill arch (A) with its associated lymphoid tissues (B). Illustrations made by Ella Maru studio.

**Figure S3.**
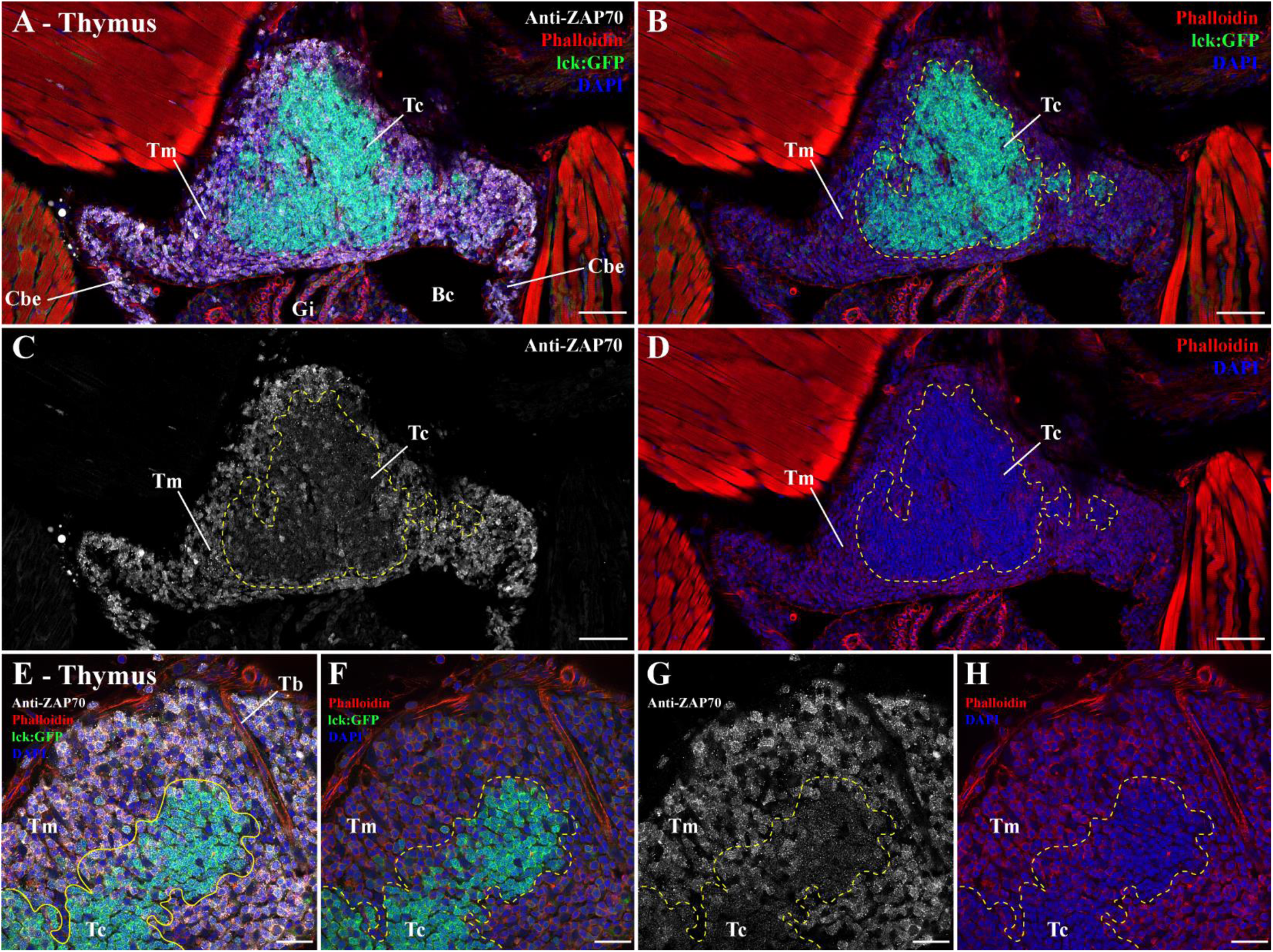
Zebrafish thymus anti-ZAP70 labeling. (A-D) Cryosections from lck:EGFP adult zebrafish, in which T cells are fluorescent (green), labeled with anti-ZAP70 antibody (white). As expected, the thymus and its GFP-positive cells are labeled by the anti-ZAP70 labeling. In the thymus cortex thymocytes are intensely packed, highly express the gene lck and display a low anti-ZAP70 labeling. In contrast, the more developed thymocytes that populate the thymus medulla showed a low lck gene expression and high anti-ZAP70 labeling. This distinction is even more striking at higher magnification (E-H). Annotations: Bc, Branchial cavity; Cbe, Cavobranchial epithelium; Gi, Gills; Tb, Trabecula; Tc, Thymus cortex and Tm, Thymus medulla. Scale bars: 50 μm (A-D) and 20 μm (E-H).

**Figure S4.**
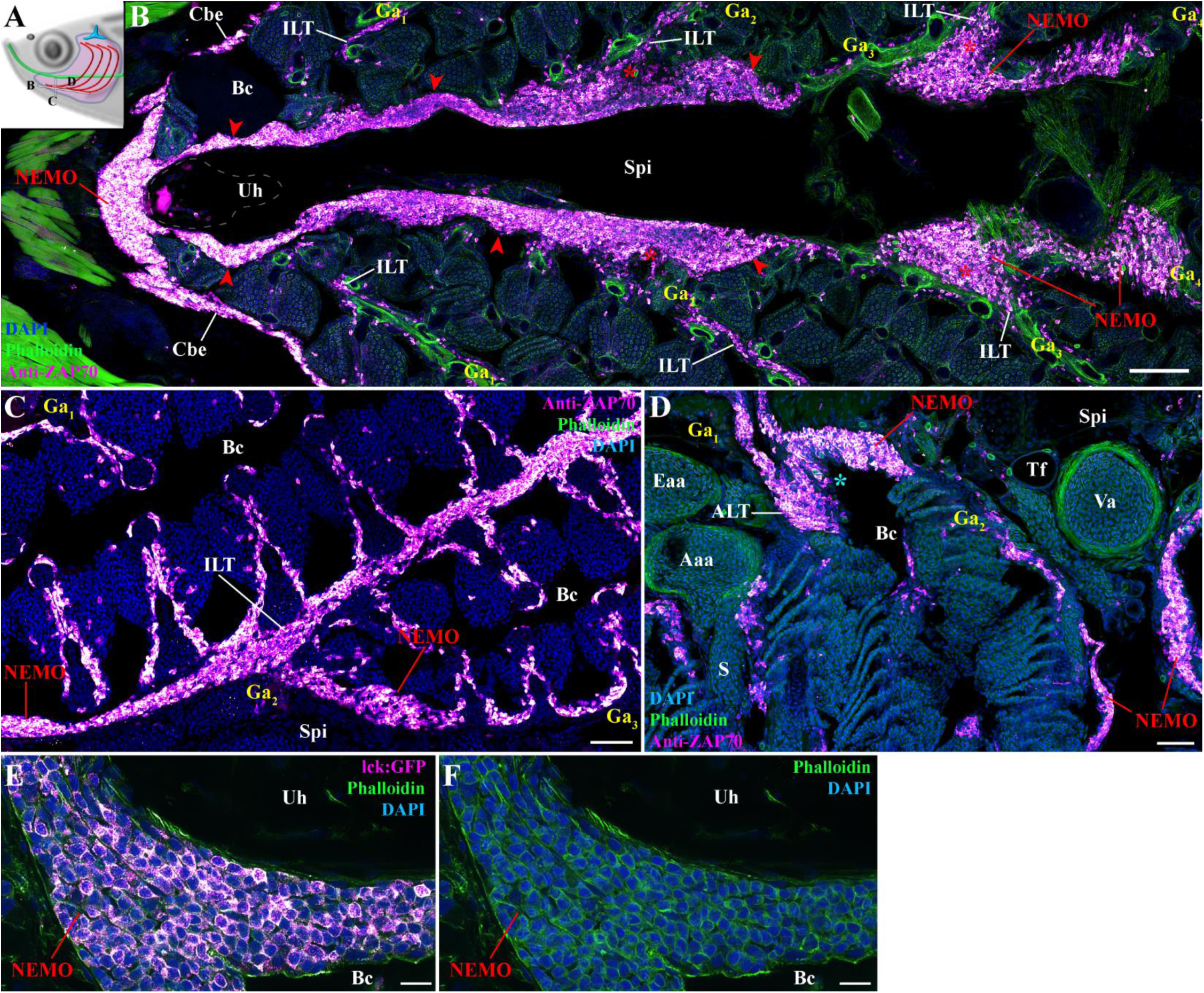
Additional information on NEMO’s identification. (A) Scheme localizing the section planes of images (B-D). (B) Additional 3D multi-field of view image of an adult zebrafish NEMO from a branchial cavity coronal cryosection labeled with anti-ZAP70 (magenta hot). The structure corresponding to NEMO is highlighted by red arrowheads. Connection sites between NEMO and ILTs are marked by red stars. (C) Additional image illustrating the continuity between NEMO and an interbranchial lymphoid tissue. (D) Additional image illustration the connection between NEMO and an amphibranchial lymphoid tissue (cyan star). (E-F) NEMO cryosection from a lck:EGFP adult zebrafish, in which T cells are fluorescent (magenta hot). Annotations: Aaa, Afferent arch artery; ALT, Amphibranchial lymphoid tissue; Bc, Branchial cavity; Cbe, Cavobranchial epithelium; Eaa, Efferent arch artery; Ga, Gill arch; ILT, Interbranchial lymphoid tissue; S, Septum; Spi; Sub-pharyngeal isthmus; Tf, Thyroid follicle; Uh, Urohyal bone and Va, Ventral aorta. Scale bars: 100 μm (B), 50 μm (C,D), and 10 μm (E,F).

**Figure S5.**
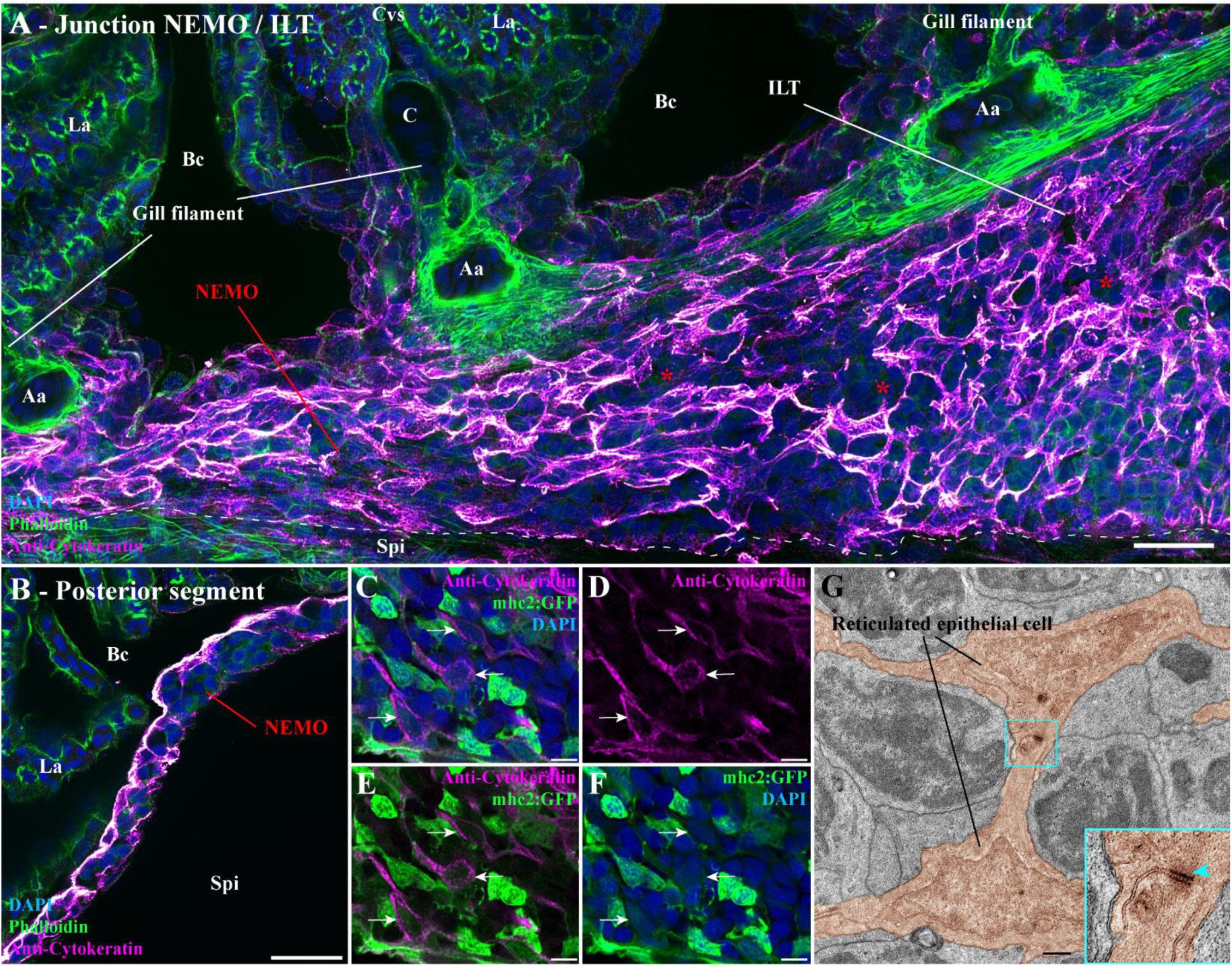
Additional information on NEMO’s network of reticulated epithelial cells. (A) Adult zebrafish cryosections labeling with anti-cytokeratin (magenta hot) display the connection site of NEMO with an ILT (red stars). (B) Network of reticulated epithelial cells at the posterior end of NEMO. (C-F) Cryosection from a mhc2:GFP adult zebrafish, in which mhc2-expressing cells are fluorescent (green), labeled with anti-cytokeratin (magenta hot). NEMO reticulated epithelial cells displayed a low mhc2 expression (white arrows). (G) Zoomed transmission electron micrograph from the ultrastructure map of Figure 2 highlighting the presence of an hemidesmosome (cyan arrowhead) at the junction of two reticulated epithelial cells (orange). Annotations: Aa, Afferent artery; Bc, Branchial cavity; C, Cartilage; ILT, Interbranchial lymphoid tissue; La, Lamellae; Spi, Sub Pharyngeal isthmus. Scale bars: 20 μm (A,B), 5 μm (C-F), and 500 nm (G).

**Figure S6.**
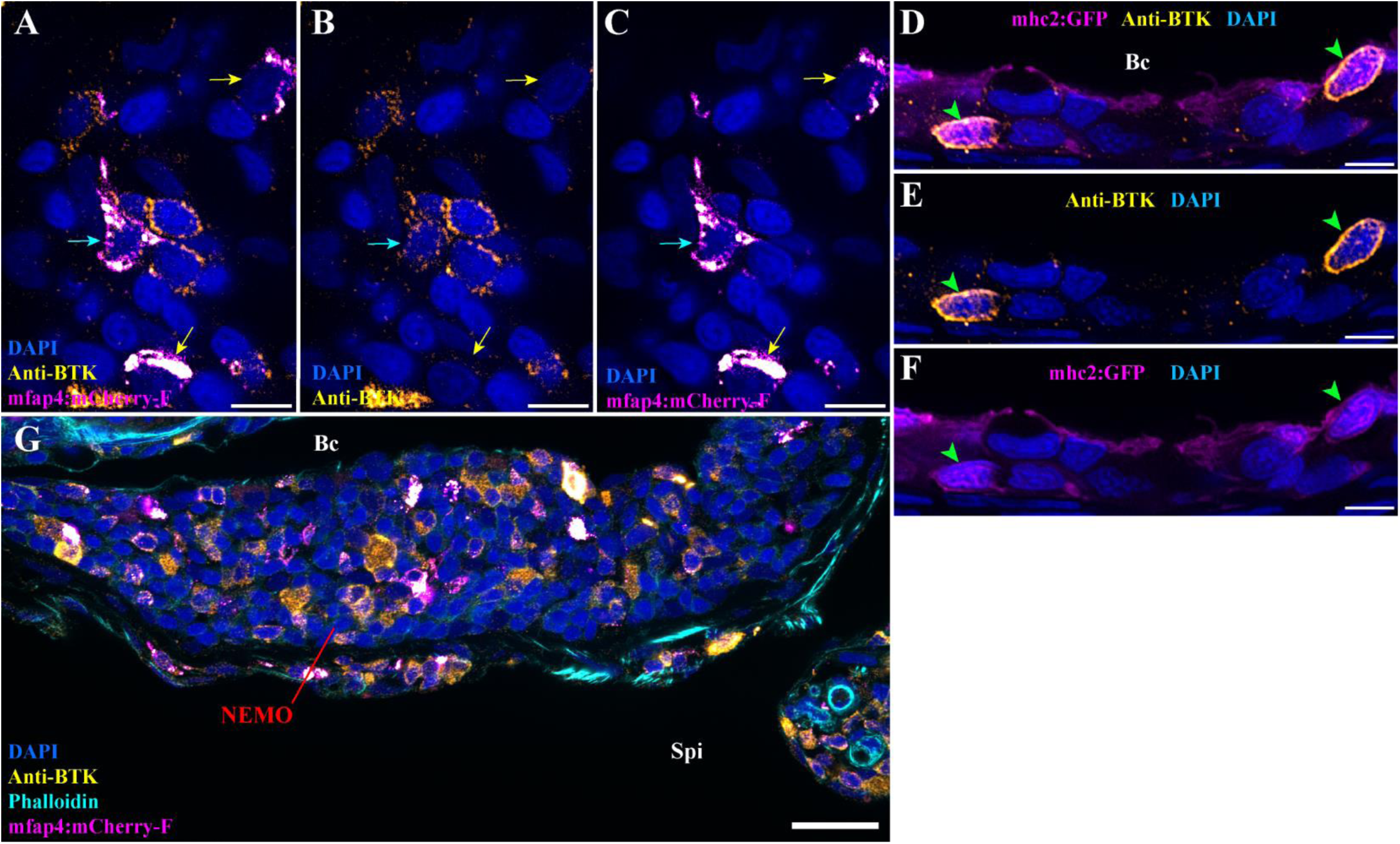
Additional information anti-BTK antibody labeling. (A-C) Anti-BTK labeling (orange hot) on mfap4:mCherry-F adult zebrafish cryosections, in which macrophage are fluorescent (magenta hot). Within NEMO, both BTK-positive (cyan arrows) and BTK-negative (yellow arrows) macrophages are observed. (D-F) Cryosection from an mhc2:GFP adult zebrafish NEMO, in which IgM expressing B cells are fluorescent (magenta hot), labeled with anti-BTK (orange hot). As expected, cells expressing IgM are also BTK-positive (green arrowheads). (G) Additional image displaying anti-BTK labeling in NEMO of a mfap4:mCherry-F adult zebrafish. Annotations: Bc, Branchial cavity and Spi, Sub-pharyngeal isthmus. Scale bars: 20 μm (G), and 5 μm (A-F).

**Figure S7.**
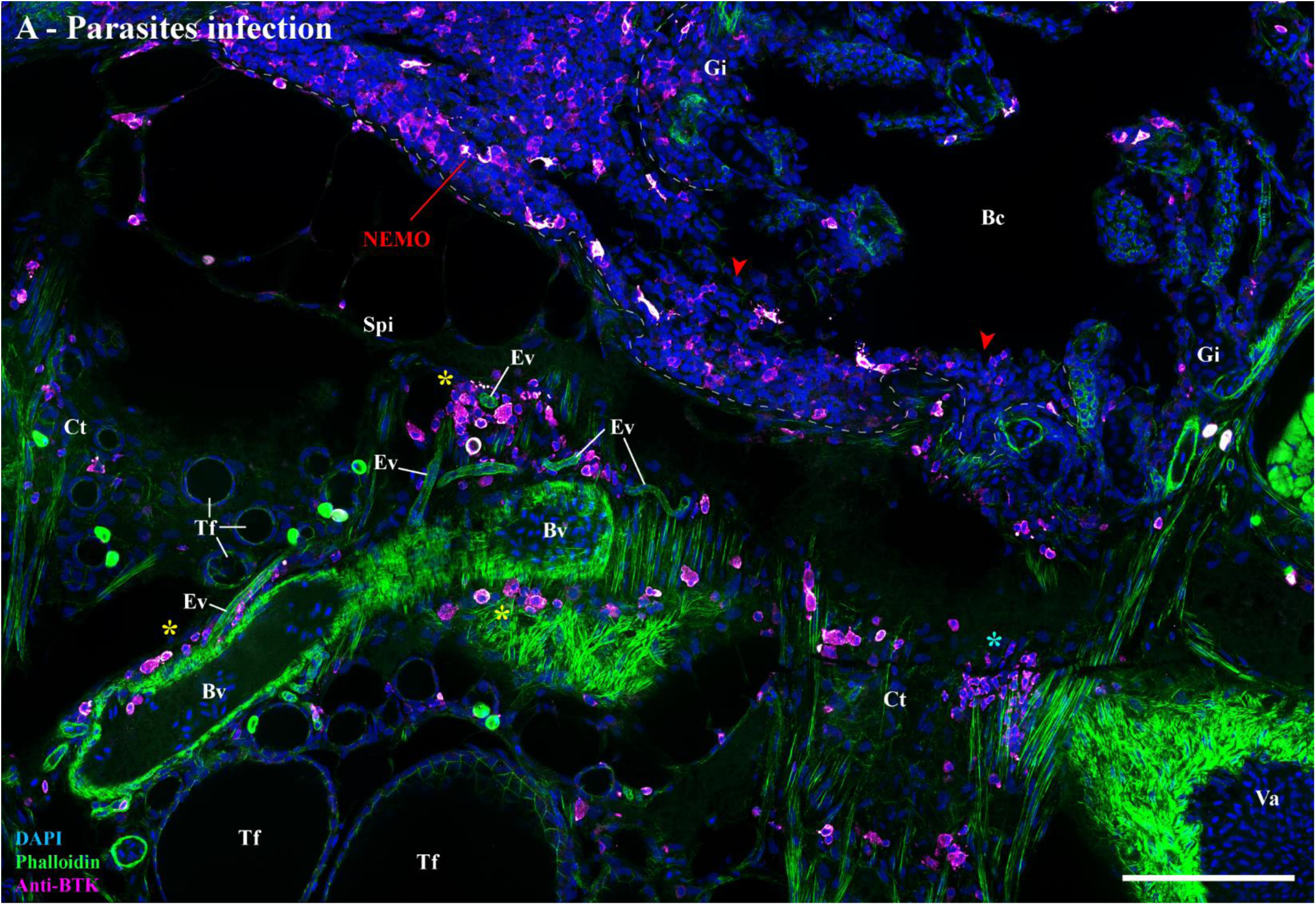
Putative plasma/B cells clusters in parasites-infected adult zebrafish. (A) Cryosection from an adult zebrafish naturally co-infected with Pseudoloma neurophilia, Pseudocapillaria tomentosa, and Myxidium streisingeri, stained with phalloidin (green) and DAPI (blue), and labeled with anti-BTK antibody (magenta hot). In addition to putative BTK-positive plasma/B cells in NEMO (red arrowheads), significant clusters of labeled cells were observed within the connective tissue (cyan star) and associated to endothelial vessels (yellow stars) of the sub-pharyngeal isthmus. Annotations: Bc, Branchial cavity; Bv, Blood vessel; Ct, Connective tissue; Ev, Endothelial vessel; Gi, Gills; Spi, Sub-pharyngeal isthmus; Tf, Thyroid follicle and Va, Ventral aorta. Scale bar: 100 μm.

**Figure S8.**
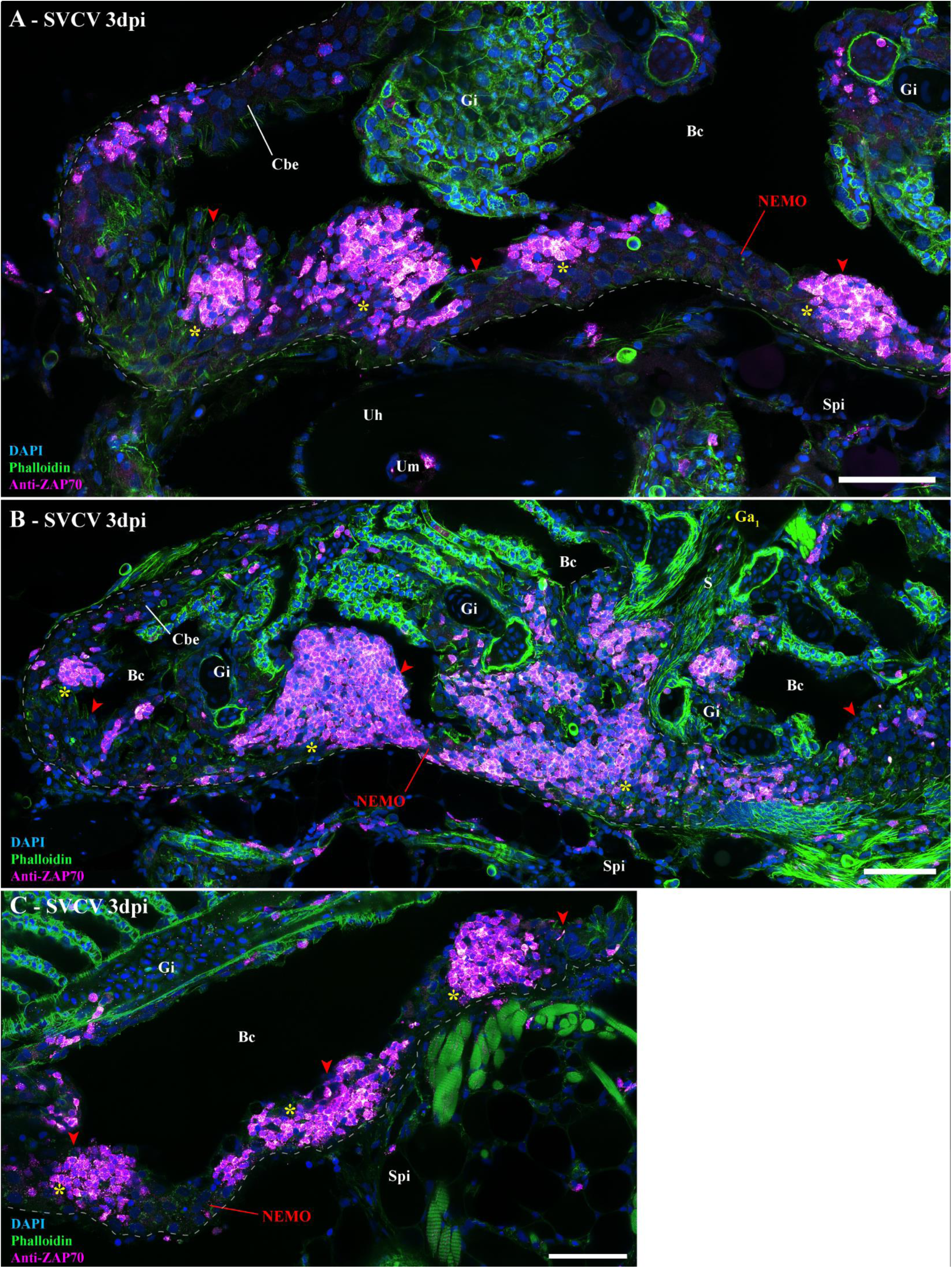
Additional images on 3 days post-SVCV infection. (A-C) Cryosections from adult zebrafish three days after a 24h bath-infection with SVCV, stained with phalloidin (green) and DAPI (blue), and labeled with anti ZAP70 (magenta hot). NEMO (red arrowheads) displayed striking aggregation of T/NK cells into distinct clusters (yellow stars). Annotations: Bc, Branchial cavity; Cbe, Cavobranchial epithelium; dpi, day post-infection; Gi, Gills; Spi, Sub-pharyngeal epithelium; SVCV, Spring viremia of carp virus; Uh, Urohyal bone and Um, Urohyal marrow. Scale bars: 50 μm (A-C).

**Figure S9.**
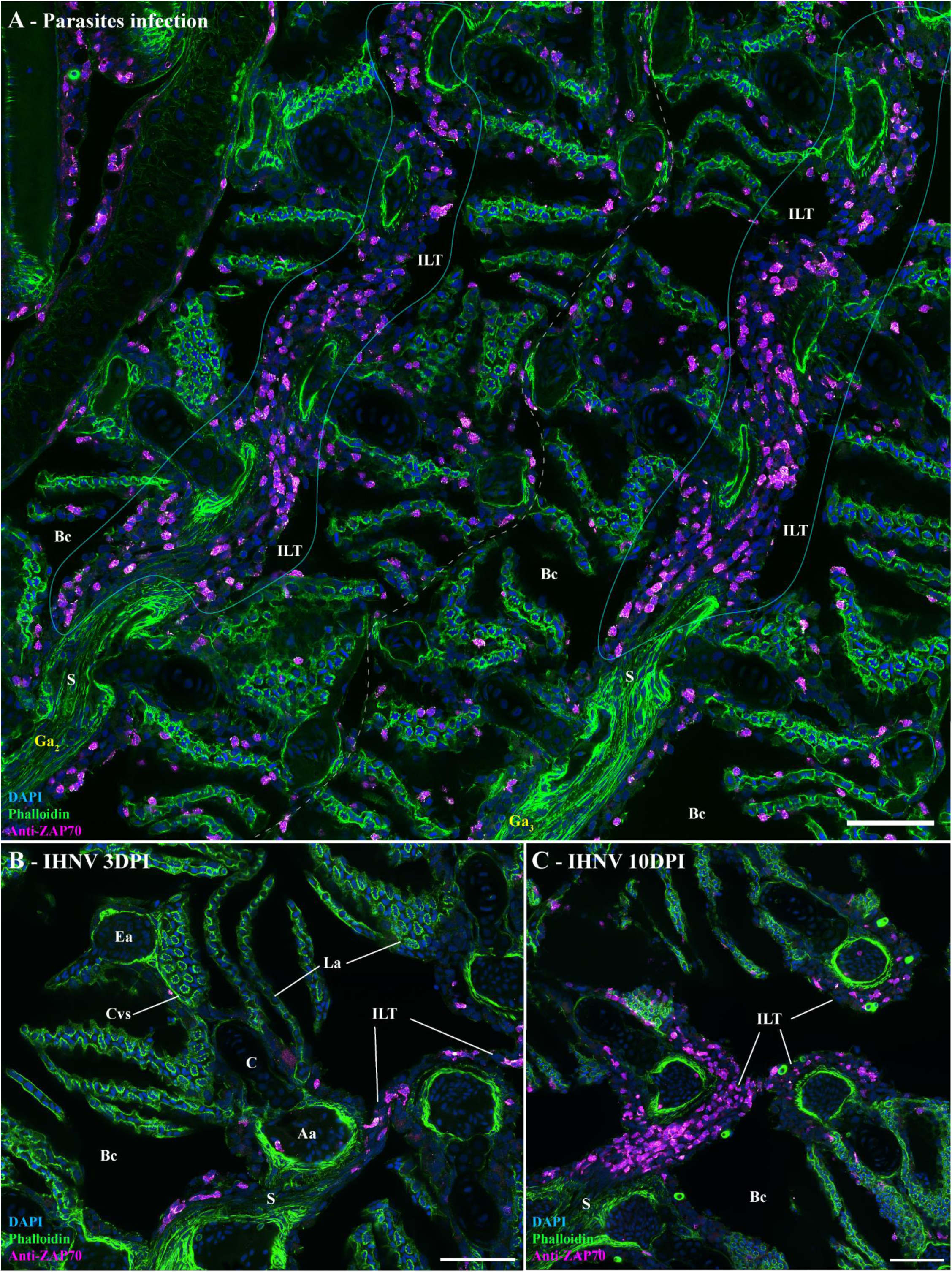
Structural response of ILT to viral and parasitic infections. (A) Cryosections displaying the interbranchial lymphoid tissue of adult zebrafish naturally co-infected with three parasitic diseases (*Pseudoloma neurophilia*, *Pseudocapillaria tomentosa*, and *Myxidium streisingeri*) stained with phalloidin (green) and DAPI (blue), and labeled with anti-ZAP70 antibody (magenta hot). The distribution of ZAP70-positive cells is more scattered than in uninfected fish and displayed small clusters of labeled cells. (B,C) Cryosection displaying the ILT of an adult zebrafish 3 days (B) and 10 days (C) following a 24h bath-infection with IHNV. Although ILTs are severely depleted at 3 dpi, they appeared replenished at 10 dpi. Annotations: Aa, Afferent artery; Bc, Branchial cavity; C, Cartilage; Cvs, Central venous sinus; Ea, Efferent artery; Ga, Gill arch; ILT, Interbranchial lymphoid tissue; La, Lamellae and S, Septum. Scale bars: 50 μm (A-C).

**Figure S10.**
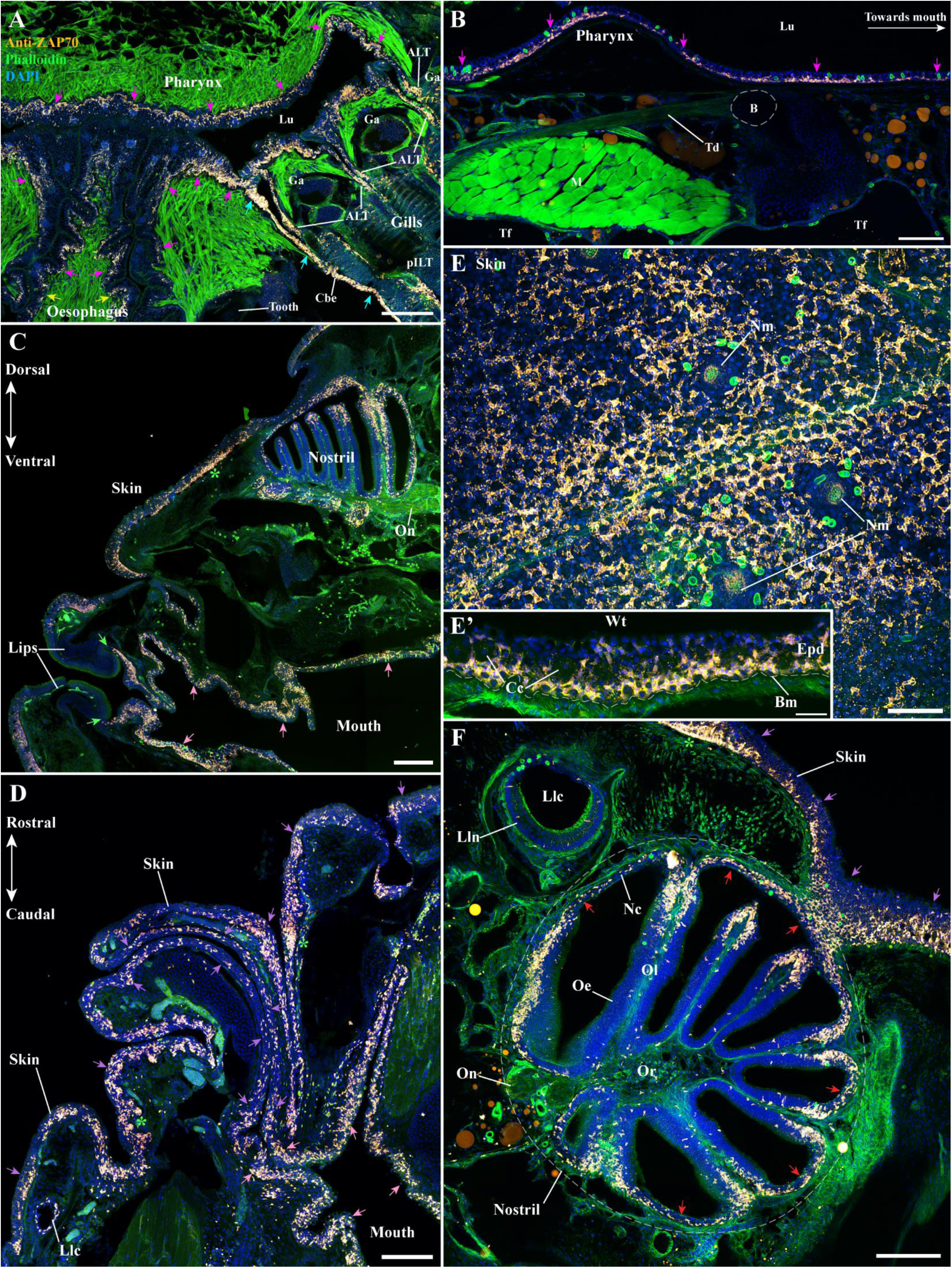
Extension of NEMO’s lymphoid network beyond the branchial cavity. (A-F) Cryosection from adult zebrafish stained with phalloidin (green) and DAPI (blue), and labeled with anti-ZAP70 (orange hot). (A) The lymphoid network of the branchial cavity, and which include NEMO, is connected to the pharynx (magenta arrows) and oesophagus (yellow arrows) via T/NK cell-rich segments of the cavobranchial epithelium (cyan arrows). This lymphoid network is observed along the length of the pharynx (B – magenta arrows) and the mouth (C,D – pink arrows). Where it is absent from the keratinized lips of the fish (C – green arrows), it connected to the skin-associated lymphoid tissue (SALT) by the sides of the mouth opening (D - purple arrows). (E) Wholemount skin of a zebrafish head labeled with anti-ZAP70 and observed from above revealed that the SALT is composed of a vast network of T/NK cells that are located at the basal layer of the epidermis and between club cells (E’) interspersed by multiple clusters of ZAP70-positive cells (C,D,F – green stars). (F) Via the organization of the SALT of the scale-less skin of the head, the lymphoid network observed in the branchial cavity is also continuous with the nasal-associated lymphoid tissue (NALT) (F – red arrows). Annotations: ALT, Amphibranchial lymphoid tissue; B, Bone; Bm, Basement membrane; Cbe, Cavobranchial epithelium; Cc, Club cells; Epd, Epidermis; Ga, Gill arch; Llc, Lateral line canal; Lln, Lateral line neuromast; Lu, Lumen; pILT, proximal Interbranchial lymphoid tissue; M, Muscles; Nc, Nasal cavity; Nm, Neuromast; Oe, Olfactory epithelium; Ol, Olfactory lamella; On, Olfactory nerve; Td, tendon; Tf, Thyroid follicle and Wt, Water. Scale bars: 200 μm (A,C), 150 μm (D,F), 100 μm (B), 50 μm (E), and 30 μm (E’).

**Video S1** – ***3D reconstruction: zebrafish NEMO.*** Reconstruction of NEMO 3D structure using serial confocal tomography on a 15 wpf zebrafish head.

**Video S2** – ***3D reconstruction: zebrafish branchial cavity region.*** Video displaying NEMO (magenta), the ventral end of gill arches (green), the ALTs (cyan), and the thymus lobes (blue) that have been 3D reconstructed using serial confocal tomography on 15 wpf zebrafish head.

**Video S3** – ***3D reconstruction: Sub-pharyngeal region of a zebrafish branchial cavity.***

Video displaying NEMO (magenta), the ventral end of gill arches (green), the ALTs (cyan), and the thymus lobes (blue) that have been 3D reconstructed using serial confocal tomography on 15 wpf zebrafish head. A section plane has been included to highlight the sub-pharyngeal region located at the convergence of the gill arches.

**Video S4** – ***3D reconstruction: Localization of NEMO, ALTs, and thymus within a zebrafish head.*** Video displaying NEMO (magenta), the ALTs (cyan), and the thymus lobes (blue) within the head of a 15 wpf zebrafish labeled with phalloidin (green).

**Video S5** – ***3D reconstruction: NEMO reticulated epithelial cell network.*** Reconstruction of a NEMO reticulated epithelial cells network from a cryosection labeled with anti-cytokeratin (red) and DAPI (blue).

**Video S6** – ***3D image: Endothelial vessels around zebrafish NEMO.*** 3D image from a fli:GFP zebrafish cryosections, in which endothelial cells are fluorescent (green), stained with phalloidin (red) and DAPI (blue), and labeled with anti-ZAP70 (white). The video display an anterior region of NEMO wrapped by endothelial vessels.

**Video S7** – ***3D reconstruction: shared reticulated epithelial cell network between NEMO and ILT.*** Reconstruction of a NEMO reticulated epithelial cells network from the cryosection labeled with anti-cytokeratin (red) and DAPI (blue) presented **Fig.S3 A**.

**Video S8** – ***3D image: Network of T/NK cells within the scale-less skin of a zebrafish head.*** The video displays the optical sections of a 3D image of the skin covering an adult zebrafish head. The acquisition was obtained from a wholemount head of zebrafish stained with phalloidin (green), DAPI (blue), and labeled with anti-ZAP70 (red hot). The optical sections are seen going from the exterior to the interior of the fish.

Supplementary videos are available on figshare: DOI: 10.6084/m9.figshare.22259698 (https://figshare.com/s/579f756a92ab85922264)

## Notes

https://figshare.com/s/579f756a92ab85922264

https://wohlmann.github.io/2019019_004_M1c/

https://wohlmann.github.io/2019019_004_N2/

### Competing Interest Statement

The authors have declared no competing interest.

### Summary of Updates

The text has been reworked and the order of figures has been changed. Scientific illustrations have been added, as well as one supplementary video.

