## Supplementary material for "Identification of a new pharyngeal mucosal lymphoid organ in zebrafish and other teleosts: tonsils in fish?"

### SUPPLEMENTARY FIGURES AND VIDEO LEGENDS


**AUTHORS**

*RESSEGUIER J [1*], NGUYEN-CHI M [2†], WOHLMANN J [3†], RIGAUDEAU D [4], SALINAS I [5], OEHLERS SH [6], WIEGERTJES GF [7], JOHANSEN FE [8], QIAO SW [9], KOPPANG EO [10], VERRIER B [11], BOUDINOT P [12‡] and GRIFFITHS G [8‡].*

[†] These authors contributed equally

[‡] These authors contributed equally

Affiliations:

[1] *Section for Physiology and Cell Biology, Departments of Biosciences and Immunology, University of Oslo, Oslo, Norway.*

[2] *LPHI, CNRS, Université de Montpellier, Montpellier, France*

[3] *Electron-Microscopy laboratory, Departments of Biosciences, University of Oslo, Oslo, Norway.*

[4] *INRAE, Université Paris-Saclay, IERP, 78350 Jouy-en-Josas, France*

[5] *Center for Evolutionary and Theoretical Immunology (CETI), Department of Biology, University of New Mexico, Albuquerque, NM, United States.*

[6] *A*STAR Infectious Diseases Labs (A*STAR ID Labs), Agency for Science, Technology and Research (A*STAR), 8A Biomedical Grove, Immunos #05-13, Singapore 138648, Singapore*

[7] *Aquaculture and Fisheries Group, Department of Animal Sciences, Wageningen University & Research, Wageningen, Netherlands*

[8] *Section for Physiology and Cell Biology, Department of Biosciences, University of Oslo, Oslo, Norway.*

[9] *Department of Immunology, Institute of Clinical Medicine, University of Oslo, Oslo, Norway.*

[10] *Unit of Anatomy, Faculty of Veterinary Medicine, Norwegian University of Life Sciences, Ås, Norway*

[11] *Laboratory of Tissue Biology and Therapeutic Engineering, UMR 5305, IBCP, CNRS, University Lyon 1, Lyon, France*

[12] *Université Paris-Saclay, INRAE, UVSQ, Virologie et Immunologie Moléculaires, Jouy-en-Josas, France.*

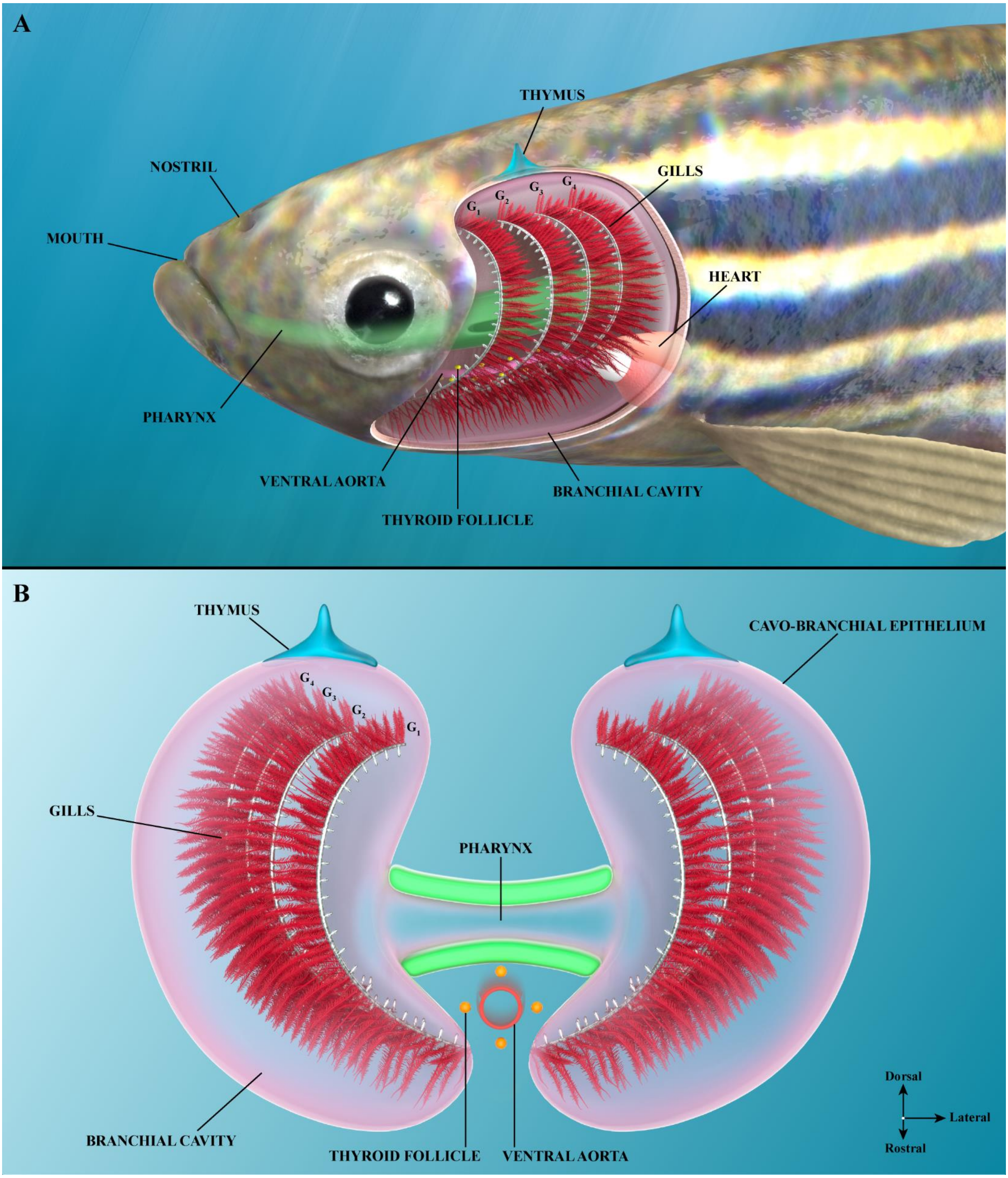


**Figure S1** – ***Organization of the adult zebrafish branchial cavity***. Illustrations of the branchial cavity tissue organization as seen from the side (A) or from a front view (B). $G_{1-4}$: First to fourth gill arch. Illustrations made by Ella Maru studio.

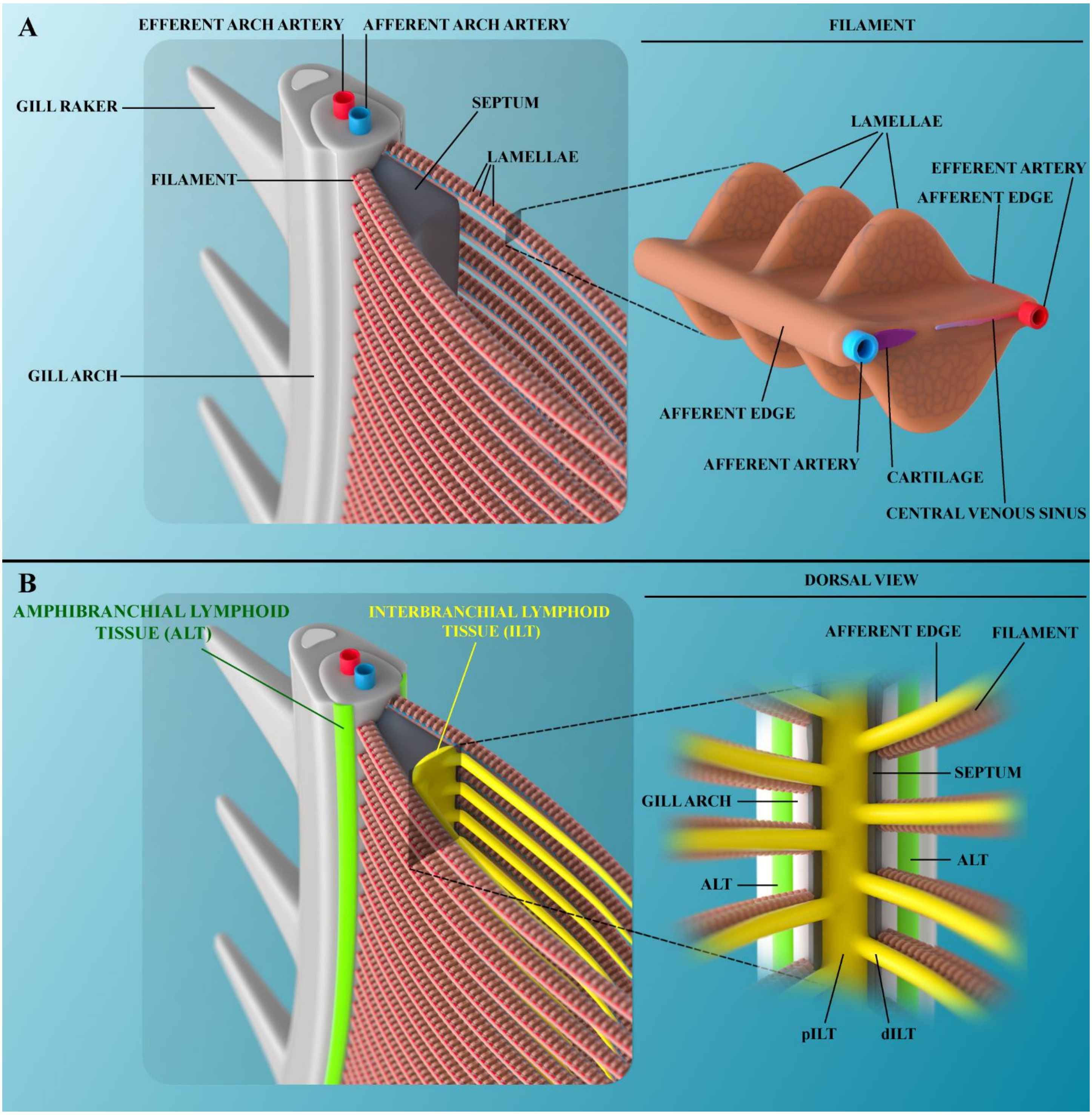


**Figure S2** – ***Organization of the adult zebrafish gills***. Illustration of a gill arch (A) with its associated lymphoid tissues (B). Illustrations made by Ella Maru studio.

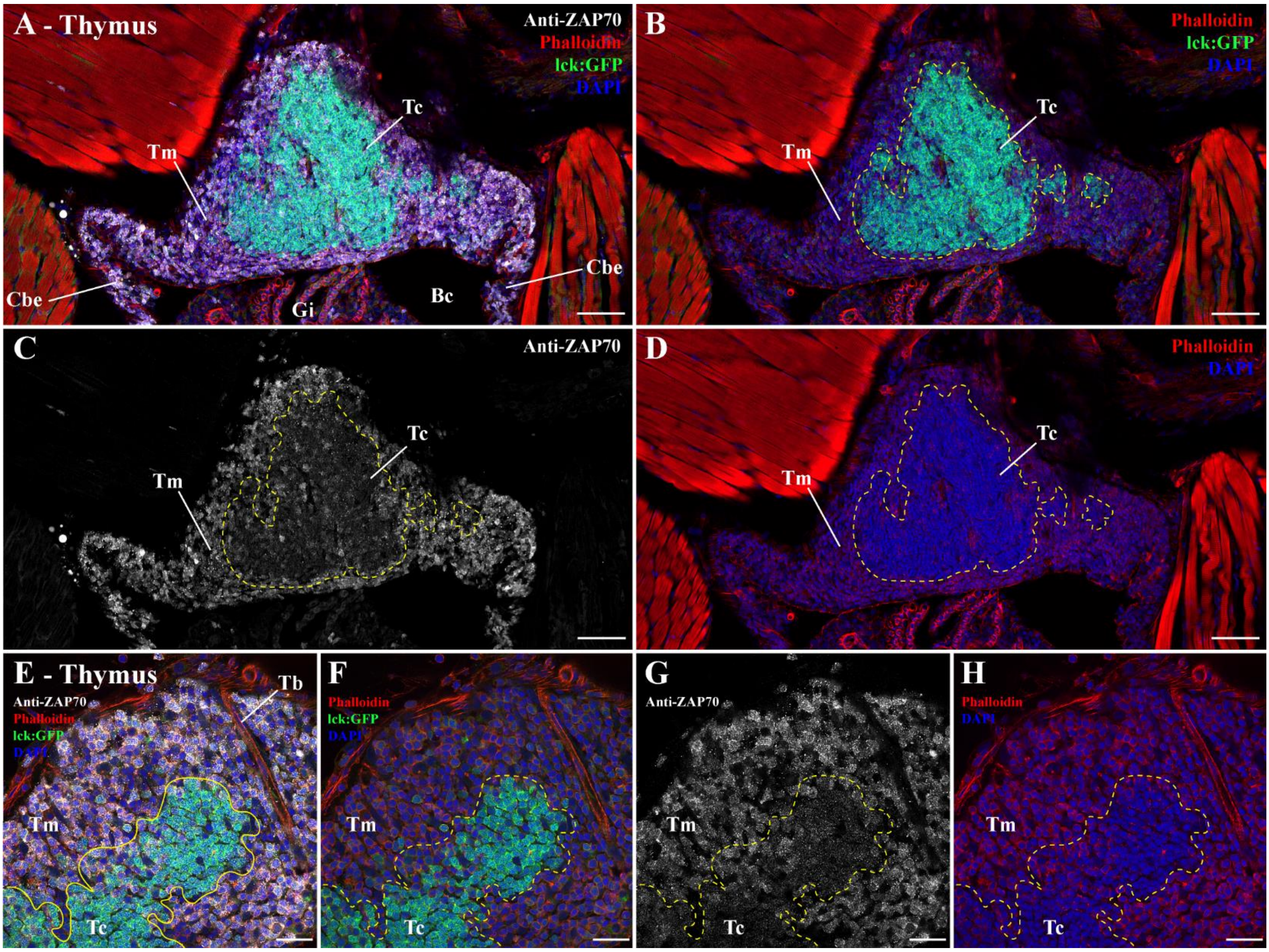


**Figure S3** – ***Zebrafish thymus anti-ZAP70 labeling***. (A-D) Cryosections from lck:EGFP adult zebrafish, in which T cells are fluorescent (green), labeled with anti-ZAP70 antibody (white). As expected, the thymus and its GFP-positive cells are labeled by the anti-ZAP70 labeling. In the thymus cortex thymocytes are intensely packed, highly express the gene lck and display a low anti-ZAP70 labeling. In contrast, the more developed thymocytes that populate the thymus medulla showed a low lck gene expression and high anti-ZAP70 labeling. This distinction is even more striking at higher magnification (E-H). Annotations: Bc, Branchial cavity; Cbe, Cavobranchial epithelium; Gi, Gills; Tb, Trabecula; Tc, Thymus cortex and Tm, Thymus medulla. Scale bars: 50 µm (A-D) and 20 µm (E-H).

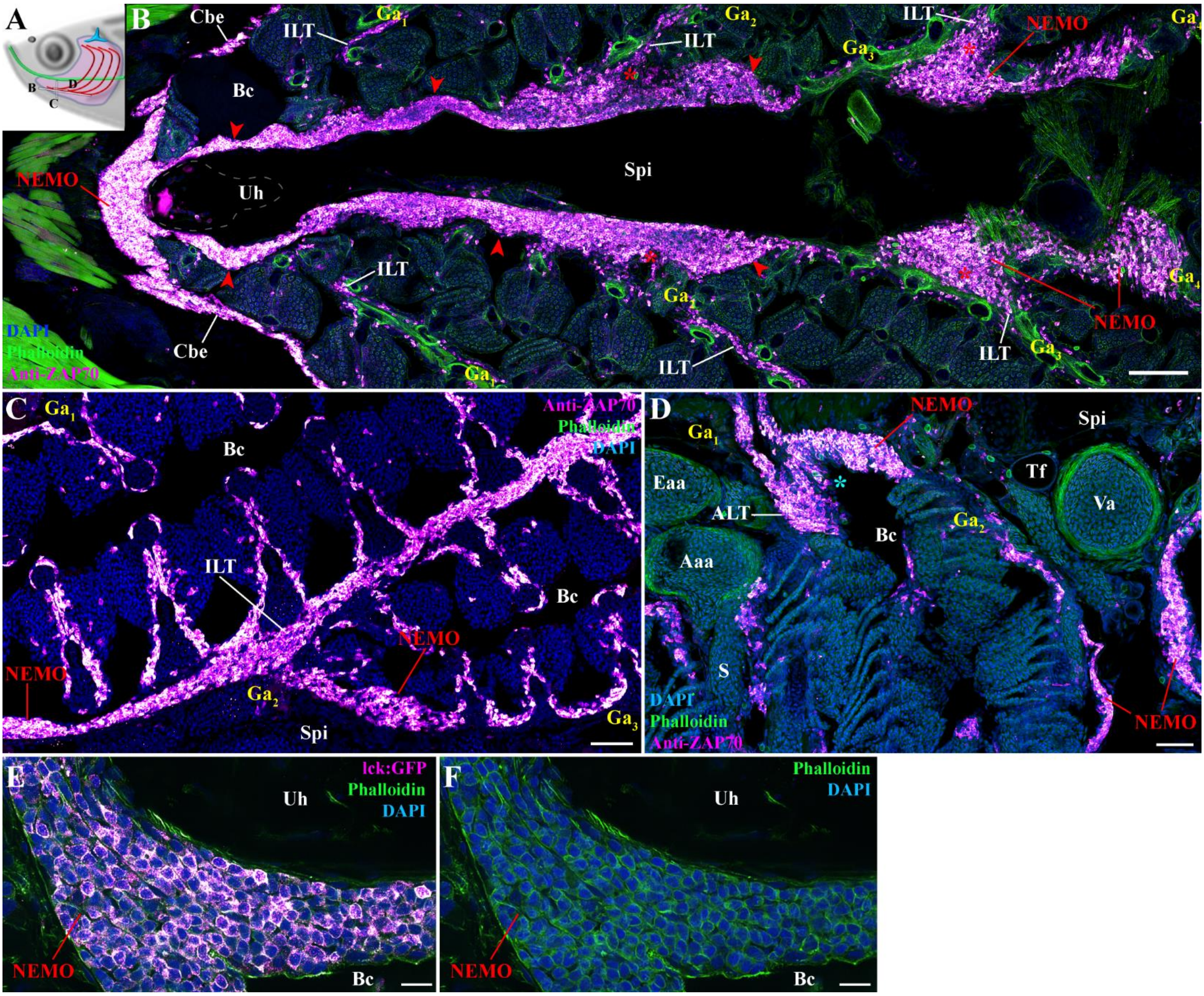


**Figure S4** – ***Additional information on NEMO's identification***. (A) Scheme localizing the section planes of images (B-D). (B) Additional 3D multi-field of view image of an adult zebrafish NEMO from a branchial cavity coronal cryosection labeled with anti-ZAP70 (magenta hot). The structure corresponding to NEMO is highlighted by red arrowheads. Connection sites between NEMO and ILTs are marked by red stars. (C) Additional image illustrating the continuity between NEMO and an interbranchial lymphoid tissue. (D) Additional image illustration the connection between NEMO and an amphibranchial lymphoid tissue (cyan star). (E-F) NEMO cryosection from a lck:EGFP adult zebrafish, in which T cells are fluorescent (magenta hot). Annotations: Aaa, Afferent arch artery; ALT, Amphibranchial lymphoid tissue; Bc, Branchial cavity; Cbe, Cavobranchial epithelium; Eaa, Efferent arch artery; Ga, Gill arch; ILT, Interbranchial lymphoid tissue; S, Septum; Spi; Sub-pharyngeal isthmus; Tf, Thyroid follicle; Uh, Urohyal bone and Va, Ventral aorta. Scale bars: 100 µm (B), 50 µm (C,D), and 10 µm (E,F).

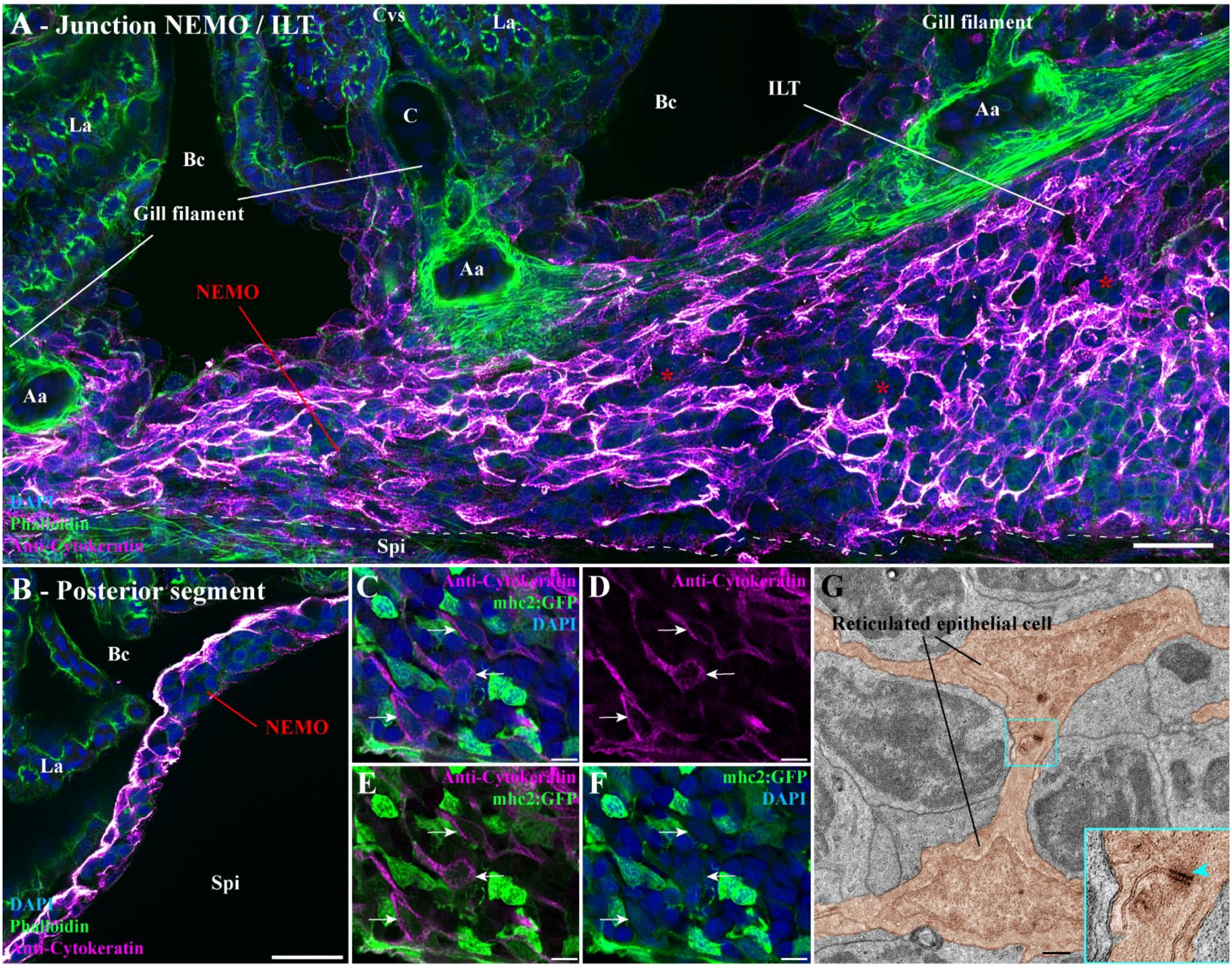


**Figure S5** – ***Additional information on NEMO's network of reticulated epithelial cells.*** (A) Adult zebrafish cryosections labeling with anti-cytokeratin (magenta hot) display the connection site of NEMO with an ILT (red stars). (B) Network of reticulated epithelial cells at the posterior end of NEMO. (C-F) Cryosection from a mhc2:GFP adult zebrafish, in which mhc2-expressing cells are fluorescent (green), labeled with anti-cytokeratin (magenta hot). NEMO reticulated epithelial cells displayed a low mhc2 expression (white arrows). (G) Zoomed transmission electron micrograph from the ultrastructure map of Figure 2 highlighting the presence of an hemidesmosome (cyan arrowhead) at the junction of two reticulated epithelial cells (orange). Annotations: Aa, Afferent artery; Bc, Branchial cavity; C, Cartilage; ILT, Interbranchial lymphoid tissue; La, Lamellae; Spi, Sub-Pharyngeal isthmus. Scale bars: 20 µm (A,B), 5 µm (C-F), and 500 nm (G).

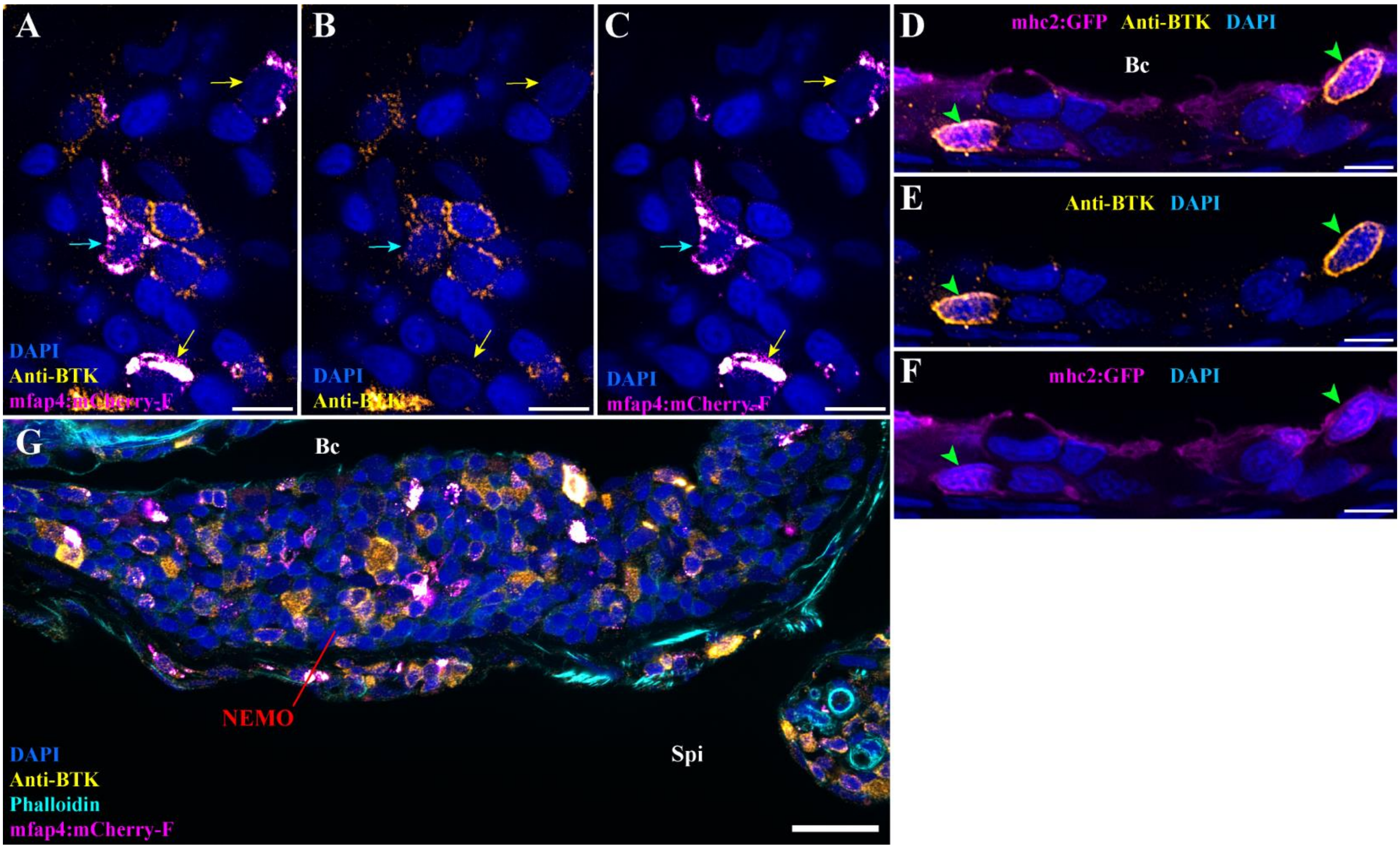


**Figure S6** – ***Additional information anti-BTK antibody labeling***. (A-C) Anti-BTK labeling (orange hot) on mfap4:mCherry-F adult zebrafish cryosections, in which macrophage are fluorescent (magenta hot). Within NEMO, both BTK-positive (cyan arrows) and BTK-negative (yellow arrows) macrophages are observed. (D-F) Cryosection from an mhc2:GFP adult zebrafish NEMO, in which IgM expressing B cells are fluorescent (magenta hot), labeled with anti-BTK (orange hot). As expected, cells expressing IgM are also BTK-positive (green arrowheads). (G) Additional image displaying anti-BTK labeling in NEMO of a mfap4:mCherry-F adult zebrafish. Annotations: Bc, Branchial cavity and Spi, Sub-pharyngeal isthmus. Scale bars: 20 µm (G), and 5 µm (A-F).

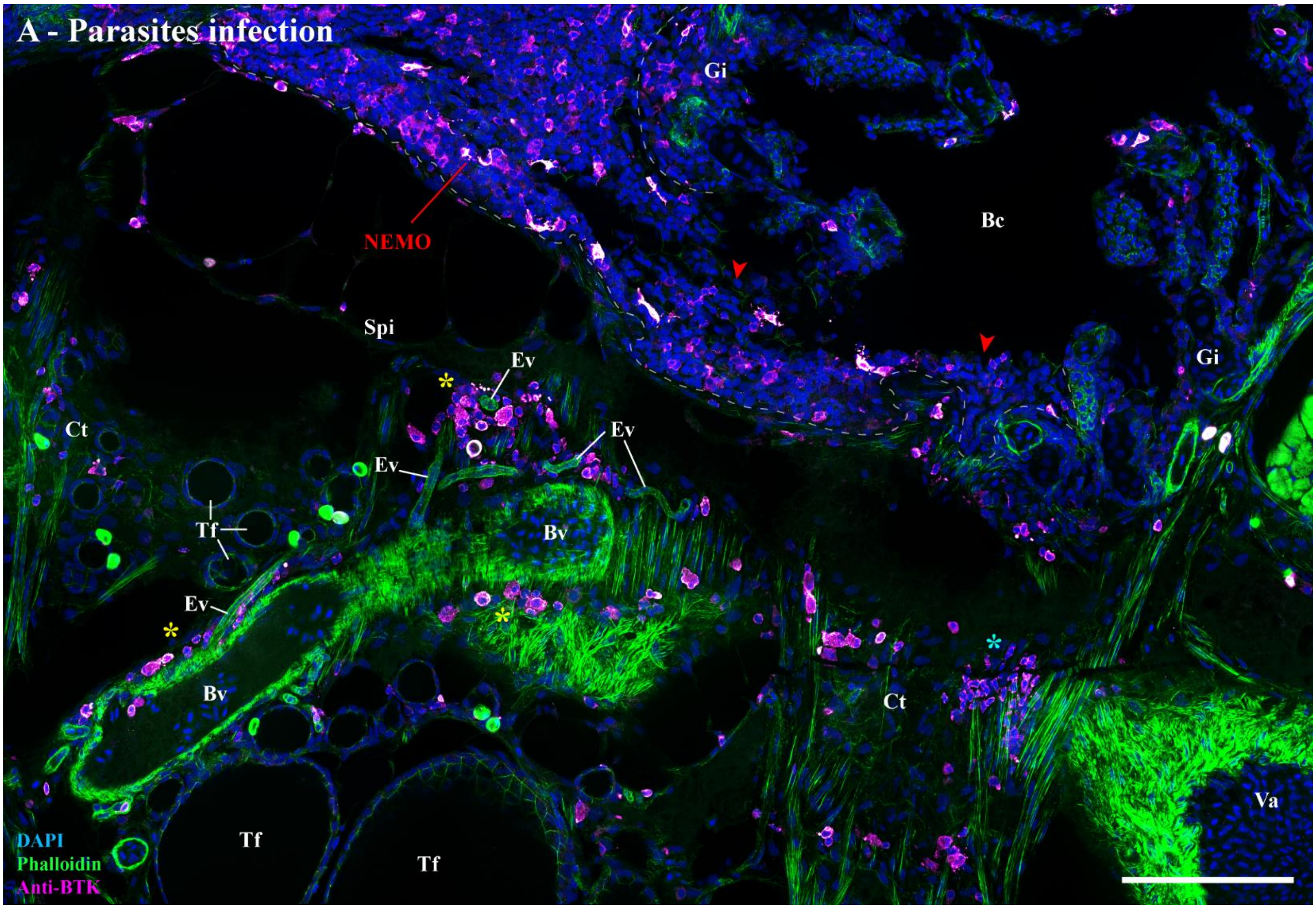


**Figure S7** – ***Putative plasma/B cells clusters in parasites-infected adult zebrafish.*** (A) Cryosection from an adult zebrafish naturally co-infected with *Pseudoloma neurophilia*, *Pseudocapillaria tomentosa*, and *Myxidium streisingeri*, stained with phalloidin (green) and DAPI (blue), and labeled with anti-BTK antibody (magenta hot). In addition to putative BTK-positive plasma/B cells in NEMO (red arrowheads), significant clusters of labeled cells were observed within the connective tissue (cyan star) and associated to endothelial vessels (yellow stars) of the sub-pharyngeal isthmus. Annotations: Bc, Branchial cavity; Bv, Blood vessel; Ct, Connective tissue; Ev, Endothelial vessel; Gi, Gills; Spi, Sub-pharyngeal isthmus; Tf, Thyroid follicle and Va, Ventral aorta. Scale bar: 100 µm.

**Figure S8** – ***Additional images on 3 days post-SVCV infection.*** (A-C) Cryosections from adult zebrafish three days after a 24h bath-infection with SVCV, stained with phalloidin (green) and DAPI (blue), and labeled with anti-ZAP70 (magenta hot). NEMO (red arrowheads) displayed striking aggregation of T/NK cells into distinct clusters (yellow stars). Annotations: Bc, Branchial cavity; Cbe, Cavobranchial epithelium; dpi, day post-infection; Gi, Gills; Spi, Sub-pharyngeal epithelium; SVCV, Spring viremia of carp virus; Uh, Urohyal bone and Um, Urohyal marrow. Scale bars: 50 µm (A-C).

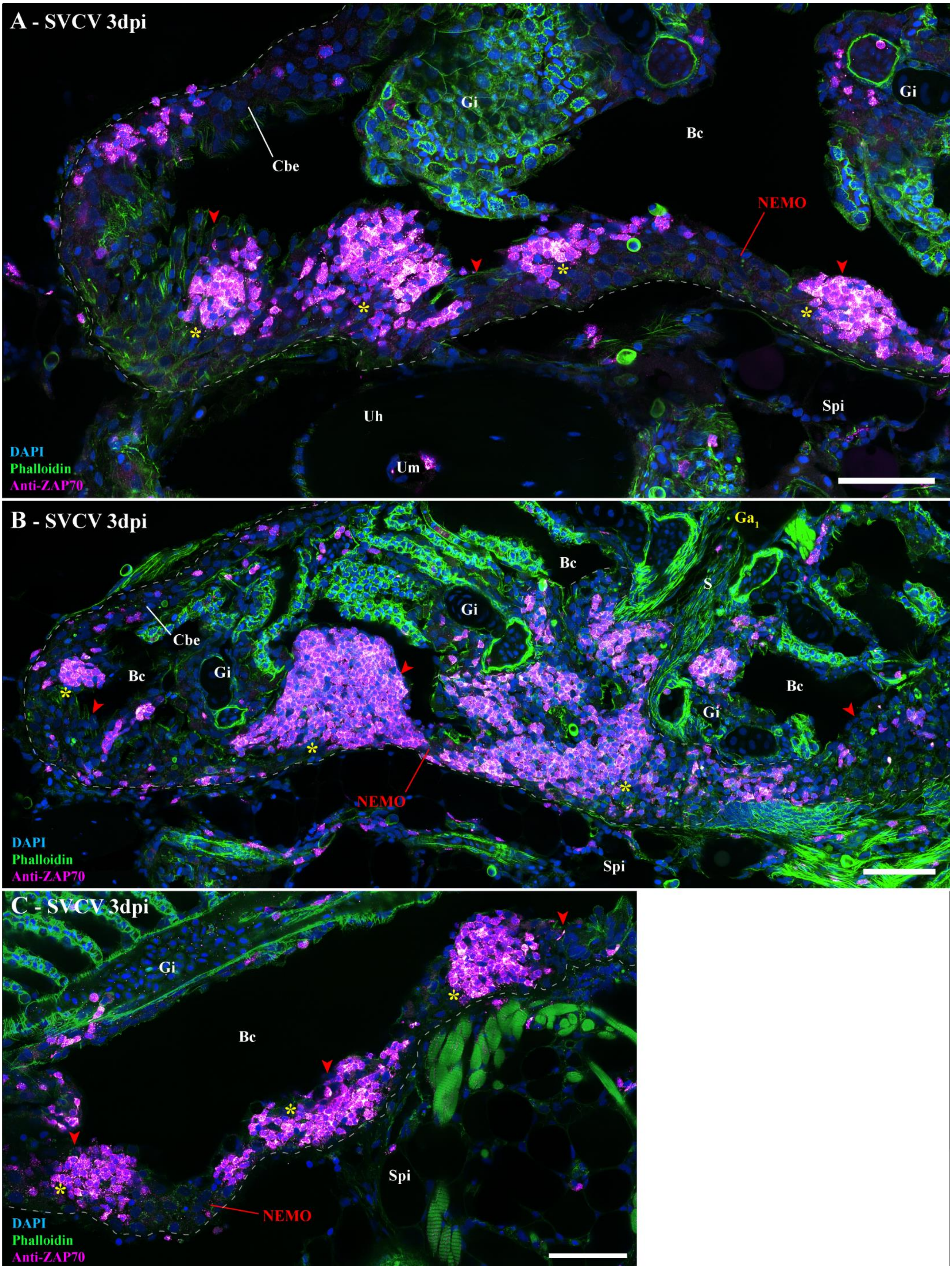
A - SVCV 3dpi
Gi
Bc
Cbe
NEMO
Uh
Um
Spi
DAPI
Phalloidin
Anti-ZAP70
B - SVCV 3dpi
Ga1
Bc
S
Gi
Cbe
Bc
Gi
NEMO
Spi
DAPI
Phalloidin
Anti-ZAP70
C - SVCV 3dpi
Gi
Bc
Spi
NEMO
DAPI
Phalloidin
Anti-ZAP70

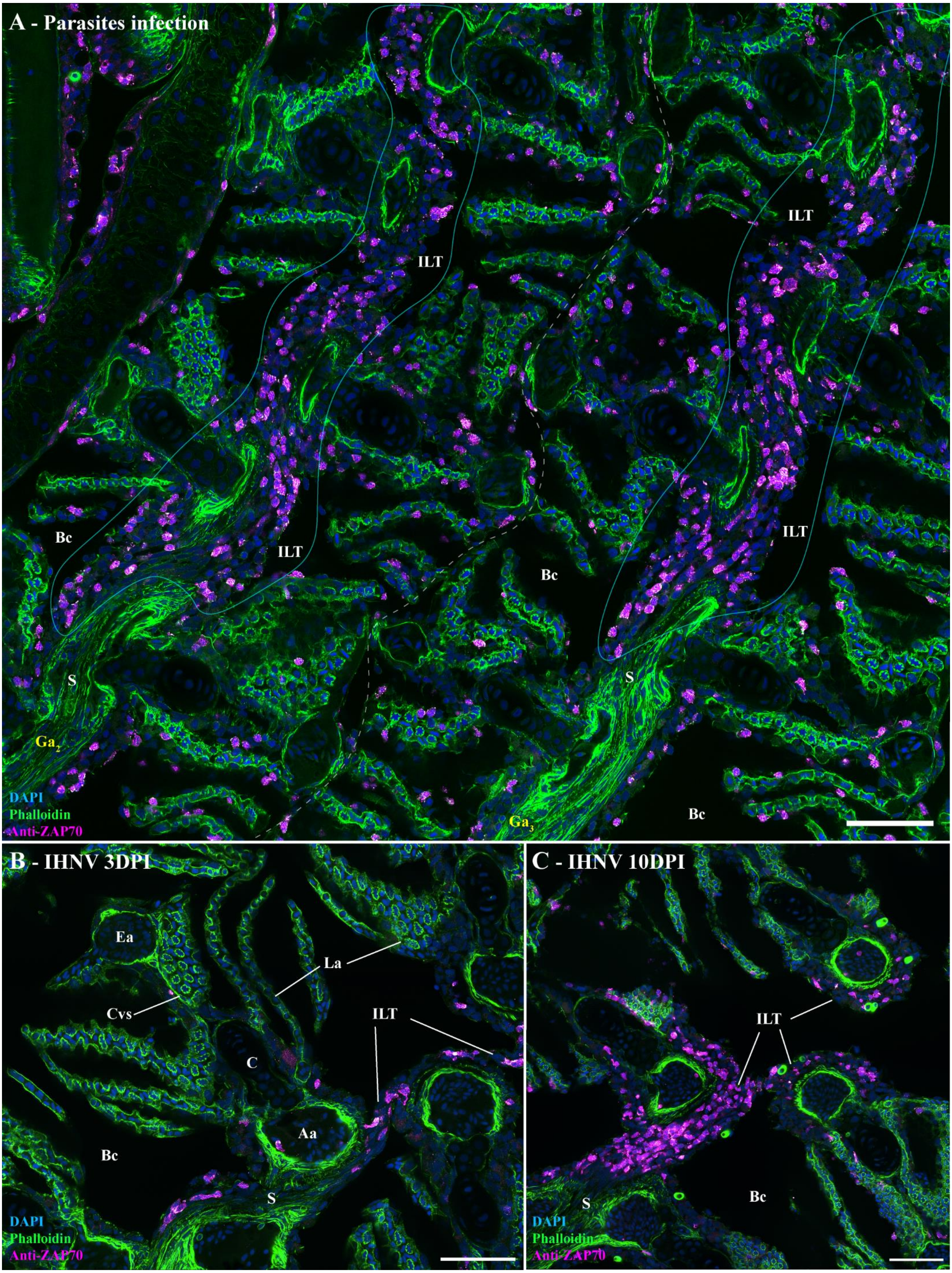
A - Parasites infection
ILT
ILT
ILT
ILT
Bc
Bc
Bc
S
S
Ga$_2$
Ga$_3$
DAPI
Phalloidin
Anti-ZAP70
B - IHNV 3DPI
Ea
La
Cvs
ILT
C
Aa
Bc
S
DAPI
Phalloidin
Anti-ZAP70
C - IHNV 10DPI
ILT
S
Bc
DAPI
Phalloidin
Anti-ZAP70

**Figure S9** – ***Structural response of ILT to viral and parasitic infections.*** (A) Cryosections displaying the interbranchial lymphoid tissue of adult zebrafish naturally co-infected with three parasitic diseases (*Pseudoloma neurophilia*, *Pseudocapillaria tomentosa*, and *Myxidium streisingeri*) stained with phalloidin (green) and DAPI (blue), and labeled with anti-ZAP70 antibody (magenta hot). The distribution of ZAP70-positive cells is more scattered than in uninfected fish and displayed small clusters of labeled cells. (B,C) Cryosection displaying the ILT of an adult zebrafish 3 days (B) and 10 days (C) following a 24h bath-infection with IHNV. Although ILTs are severely depleted at 3 dpi, they appeared replenished at 10 dpi. Annotations: Aa, Afferent artery; Bc, Branchial cavity; C, Cartilage; Cvs, Central venous sinus; Ea, Efferent artery; Ga, Gill arch; ILT, Interbranchial lymphoid tissue; La, Lamellae and S, Septum. Scale bars: 50 µm (A-C).

**Figure S10** – ***Extension of NEMO's lymphoid network beyond the branchial cavity.*** (A-F) Cryosection from adult zebrafish stained with phalloidin (green) and DAPI (blue), and labeled with anti-ZAP70 (orange hot). (A) The lymphoid network of the branchial cavity, and which include NEMO, is connected to the pharynx (magenta arrows) and oesophagus (yellow arrows) via T/NK cell-rich segments of the cavobranchial epithelium (cyan arrows). This lymphoid network is observed along the length of the pharynx (B – magenta arrows) and the mouth (C,D – pink arrows). Where it is absent from the keratinized lips of the fish (C – green arrows), it connected to the skin-associated lymphoid tissue (SALT) by the sides of the mouth opening (D - purple arrows). (E) Wholemount skin of a zebrafish head labeled with anti-ZAP70 and observed from above revealed that the SALT is composed of a vast network of T/NK cells that are located at the basal layer of the epidermis and between club cells (E') interspersed by multiple clusters of ZAP70-positive cells (C,D,F – green stars). (F) Via the organization of the SALT of the scale-less skin of the head, the lymphoid network observed in the branchial cavity is also continuous with the nasal-associated lymphoid tissue (NALT) (F – red arrows). Annotations: ALT, Amphibranchial lymphoid tissue; B, Bone; Bm, Basement membrane; Cbe, Cavobranchial epithelium; Cc, Club cells; Epd, Epidermis; Ga, Gill arch; Llc, Lateral line canal; Lln, Lateral line neuromast; Lu, Lumen; pILT, proximal Interbranchial lymphoid tissue; M, Muscles; Nc, Nasal cavity; Nm, Neuromast; Oe, Olfactory epithelium; Ol, Olfactory lamella; On, Olfactory nerve; Td, tendon; Tf, Thyroid follicle and Wt, Water. Scale bars: 200 µm (A,C), 150 µm (D,F), 100 µm (B), 50 µm (E), and 30 µm (E').

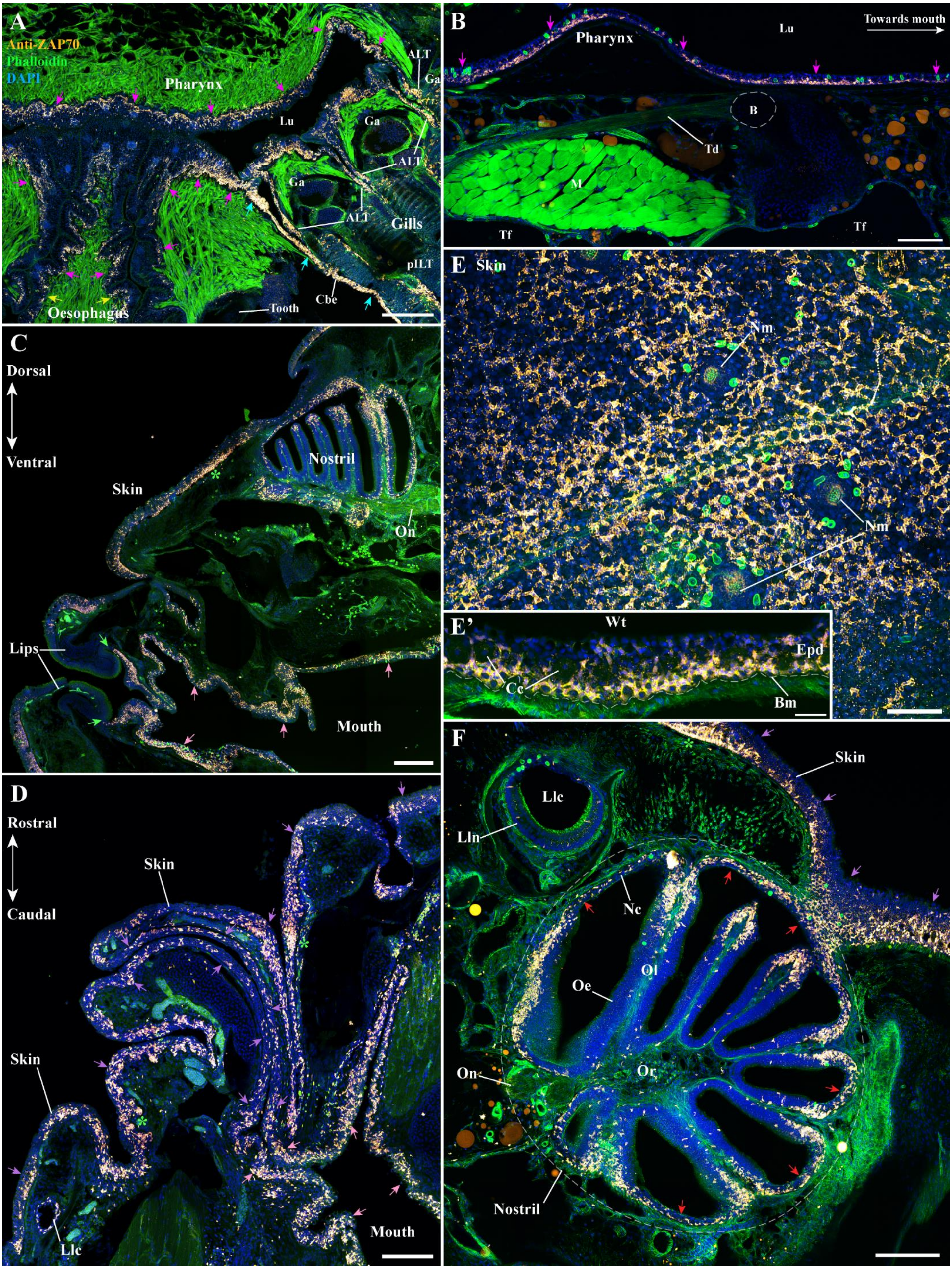
A
Anti-ZAP70
Phalloidin
DAPI
Pharynx
Lu
ALT
Ga
Gills
pILT
Cbe
Tooth
Oesophagus
B
Pharynx
Lu
Towards mouth
B
Td
M
Tf
C
Dorsal
Ventral
Skin
Nostril
On
Lips
Mouth
D
Rostral
Caudal
Skin
Llc
Mouth
E
Skin
Nm
E'
Wt
Epd
Cc
Bm
F
Skin
Llc
Lln
Nc
Ol
Oe
On
Or
Nostril

**Video S1** – ***3D reconstruction: zebrafish NEMO.*** Reconstruction of NEMO 3D structure using serial confocal tomography on a 15 wpf zebrafish head.
